# Gravin-associated kinase signaling networks coordinate γ-tubulin organization at mitotic spindle poles

**DOI:** 10.1101/2020.06.12.148171

**Authors:** Paula J. Bucko, Irvin Garcia, Ridhima Manocha, Akansha Bhat, Linda Wordeman, John D. Scott

## Abstract

Mitogenic signals that regulate cell division often proceed through multi-enzyme assemblies within defined intracellular compartments. The anchoring protein Gravin restricts the action of mitotic kinases and cell-cycle effectors to defined mitotic structures. In this report we discover that genetic deletion of Gravin disrupts proper accumulation and asymmetric distribution of γ-tubulin during mitosis. We utilize a new precision pharmacology tool, Local Kinase Inhibition (LoKI), to inhibit the Gravin binding partner polo-like kinase 1 (Plk1) at spindle poles. Using a combination of gene-editing approaches, quantitative imaging, and biochemical assays we provide evidence that disruption of local Plk1 signaling underlies the γ-tubulin distribution defects observed with Gravin loss. Our study uncovers a new role for Gravin in coordinating γ-tubulin recruitment during mitosis and illuminates the mechanism by which signaling enzymes regulate this process at a distinct subcellular location.

## Introduction

Spatial biology is an emerging aspect of biomedicine wherein investigators study how the subcellular location of enzymes underlies health and disease (1,2). At the molecular level, intracellular targeting of cell signaling enzymes is achieved through the interaction with anchoring, adaptor, or scaffolding proteins (3). A-kinase anchoring proteins (AKAPs) are a prototypic example of these signal-organizing proteins that are classified by their ability to anchor protein kinase A (PKA) ((4,5); Bucko and Scott, in press). However, numerous studies have identified other classes of protein kinases, phosphatases, small GTPases and effector proteins that are AKAP-binding partners (6). Thus, an important function of AKAPs is to provide a locus for the processing, integration, and termination of chemical stimuli by constraining these signaling molecules at defined subcellular locations. Recent findings highlight that AKAPs provide spatial and temporal synchronization of protein kinases that control the mammalian cell cycle. Several multivalent AKAPs such as pericentrin, AKAP450, and Gravin have been implicated in targeting PKA and other signaling enzymes to mitotic structures (7–10). For example, during mitosis Gravin anchors Aurora A and polo-like kinase 1 (Plk1) at the spindle poles (11). We previously showed that mitotic defects and cell-cycle delay ensue upon displacement of either enzyme from the Gravin scaffold. However, given the breadth of targets for these enzymes, a mechanistic dissection of their role in regulating mitotic machinery remains complex.

γ-tubulin participates in various aspects of cell division including centrosome duplication, chromosome segregation, and mitotic progression (12–14). Importantly, γ-tubulin also initiates nucleation of microtubules at the spindle poles during mitosis (15,16). Aberrant expression and subcellular distribution of γ-tubulin has been documented in several cancers including gliomas, medulloblastomas, breast cancer, and non-small cell lung cancer (17–21). Thus, gaining greater insight into how γ-tubulin is regulated is paramount to understanding its role in disease and for the development of novel therapeutics that target this protein (22,23).

In this study, we discover that Gravin loss impairs the accumulation of γ-tubulin at the spindle poles during mitosis. We utilize a recently developed precision pharmacology tool, Local Kinase Inhibition (LoKI), to demonstrate that targeted inhibition of centrosomal Plk1, alters the accumulation and asymmetric distribution γ-tubulin at mitotic spindle poles. Finally, we show that deletion of Gravin disrupts the formation of Nedd1/γ-tubulin sub-complexes that are necessary for tubulin ring assembly, thus illuminating a key centrosome-specific function of this anchoring protein.

## Results

### Gravin loss perturbs accumulation of γ-tubulin at mitotic spindle poles

Gravin organizes several signaling elements during mitosis (10,24,25). Conversely, removal of Gravin disrupts the localization of active kinases at mitotic centrosomes and is associated with spindle defects (11,26). To ensure the proper assembly of mitotic spindles, γ-tubulin initiates nucleation of microtubules at the spindle poles (27). We examined whether cells with genetically ablated Gravin exhibited defects in the expression or localization of this critical spindle assembly component. First, immunoblots of mouse embryonic fibroblasts (MEFs) from wildtype (WT) and Gravin null (−/−) mice established that total cellular protein expression of γ-tubulin was similar for both genotypes (Figure 1A). Next, immunofluorescence detection confirmed that γ-tubulin (magenta) decorates spindle poles in wildtype MEFs during mitosis (Figure 1B). Counterstaining with α-tubulin (green) and DAPI (blue) revealed the mitotic spindle and DNA, respectively (Figure 1B). Strikingly, γ-tubulin is much less concentrated at the spindle poles in Gravin null cells (Figure 1C). Representative heat maps highlight this phenomenon (Figure 1B-E). Likewise, surface plots depicting maximum intensity signals further illustrate this result (Figure 1D-G). Quantitative analysis using integrated intensity measurements revealed 33.4% less γ-tubulin at mitotic spindle poles in Gravin null MEFs as compared to wildtype cells (Figure 1H). Complementary experiments in HeLa cells stably expressing a control or Gravin shRNA provide further evidence for this phenomenon (Figure S1, A & B). Gravin-depleted HeLa cells display 23.6% less γ-tubulin at the spindle poles than controls (Figure S1C). Furthermore, this defect was corrected upon rescue with murine Gravin (Figure S1D-F). Collectively, these findings reveal that accumulation of γ-tubulin during mitosis is disrupted in cells lacking Gravin.

**Figure 1:**
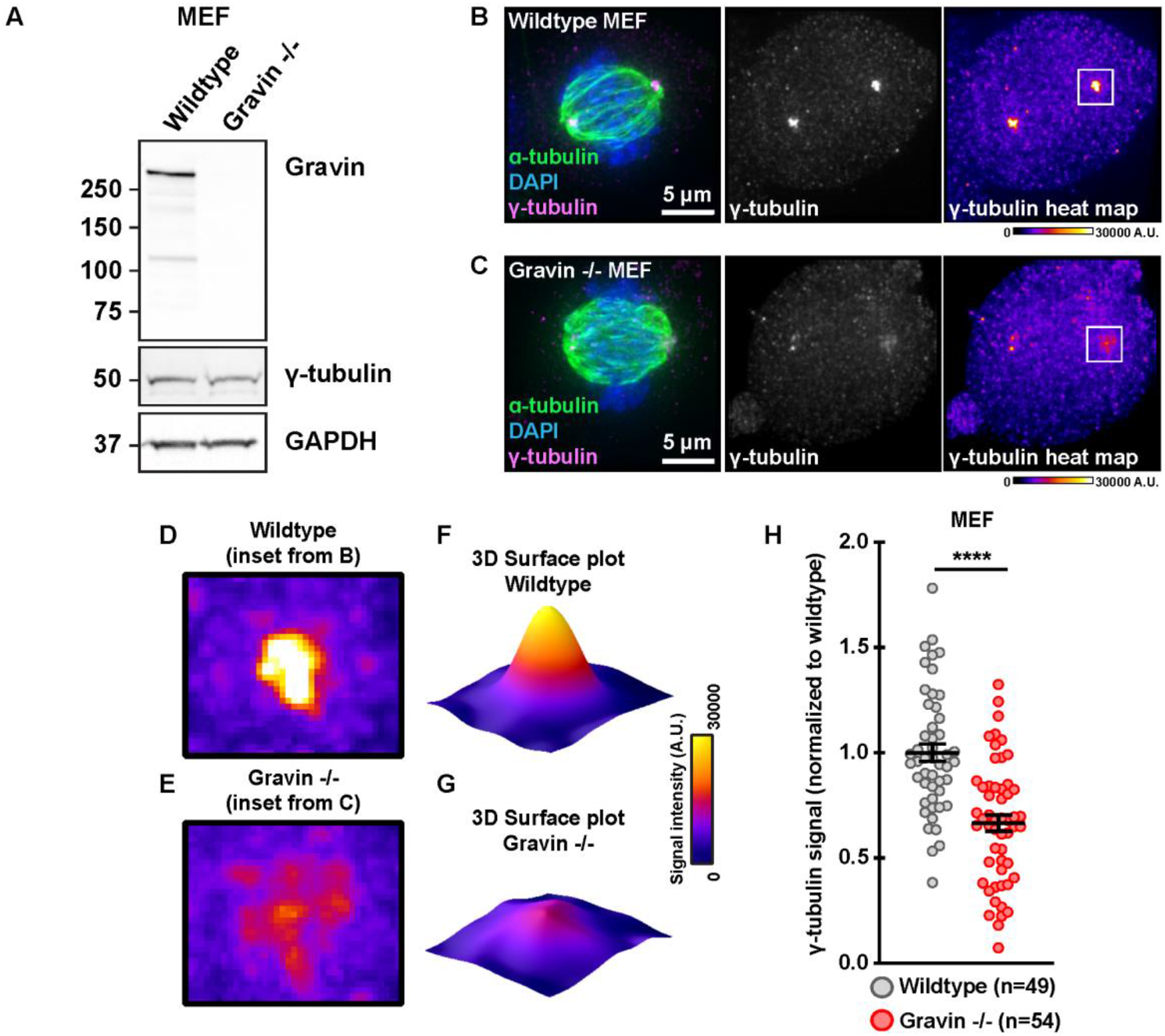
Loss of Gravin perturbs accumulation of γ-tubulin at mitotic spindle poles. **A**. Immunoblot detection of Gravin (top), γ-tubulin (middle), and GAPDH (bottom) in wildtype and Gravin null (-/-) mouse embryonic fibroblasts (MEFs). **B & C**. Immunofluorescence of representative wildtype (**B**) and Gravin -/- (**C**) mitotic cells. Composite images (left) show α-tubulin (green), DAPI (blue), and γ-tubulin (magenta). Distribution of γ-tubulin is presented in grayscale (middle) and as pseudo-color heat maps (right). Signal intensity scale (A.U.) is shown below. **D & E**. Magnified heat-maps from image insets in B and C (white boxes) highlight γ-tubulin signal in wildtype (**D**) and Gravin -/- (**E**) cells. **F, G**. Surface plots representing signal intensity of γ-tubulin in wildtype (**F**) and Gravin -/- (**G**) cells. **H**. Quantification of γ-tubulin immunofluorescence at spindle poles in wildtype (gray) and Gravin -/-(red) MEFs. Points represent individual cells (n). Data are normalized to wildtype controls; wildtype, n=49, Gravin -/-, n=54, ****p<0.0001; Experiments were conducted four times (N=4). P values were calculated by unpaired two-tailed Student’s t-test. Data are mean ± s.e.m.

### Generation and validation of genetically engineered human Gravin KO cells

While our initial findings postulate that Gravin loss disrupts γ-tubulin targeting to mitotic spindle poles, it is not clear whether complete deletion of Gravin in human cells produces a similar outcome. To generate Gravin knockout (KO) human lines we used CRISPR/Cas9-mediated genome editing to disrupt the *AKAP12* gene on chromosome 6 in U2OS osteosarcoma cells (Figure 2A). Targets were selected within the exon of *AKAP12* that is shared by all three Gravin isoforms, α, β, and γ (Figure 2A, middle schematic). We employed either a single guide RNA (gRNA) directed toward target 1 (Gravin KO) or a combination of two gRNAs directed toward both targets 1 and 2 (Gravin KO #2) to generate independent Gravin KO cell lines (Figure 2A, bottom schematic).

**Figure 2:**
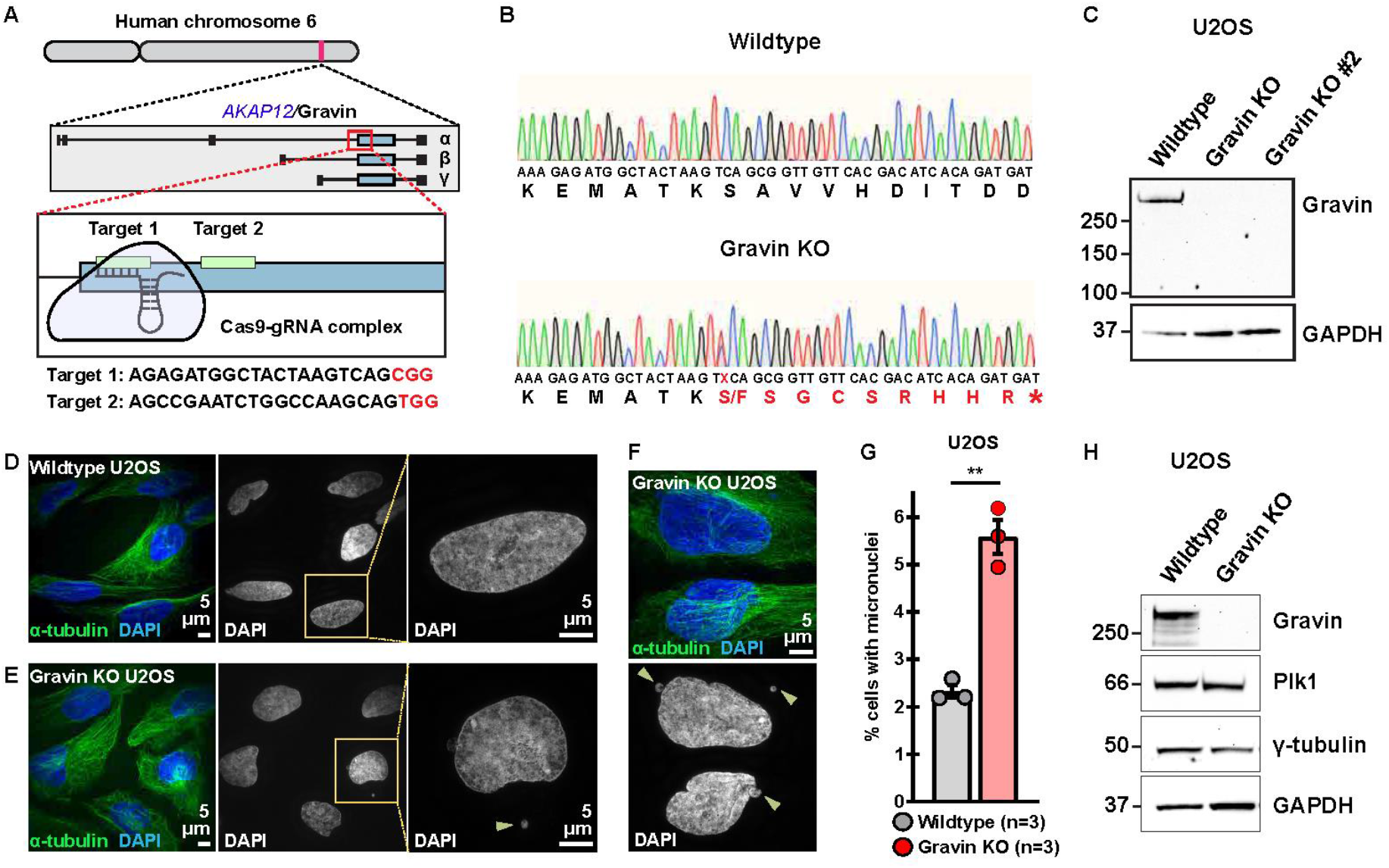
Generation of Gravin knockout U2OS cells. **A**. CRISPR-Cas9 gene editing of human chromosome six in U2OS cells to disrupt the Gravin-encoding gene, *AKAP12* (top). Targets are directed to the exon that is shared by all three Gravin isoforms, α, β, and γ (middle). Two unique guide RNAs (gRNAs) were designed to targets 1 and 2 (bottom). Target sequences are presented bellow with PAM sites highlighted in red. **B**. Traces depicting nucleic (small letters) and amino (large letters) acid sequences for a representative wildtype (top) and Gravin KO (bottom) clone. A mutation present in Gravin KO cells alters the protein coding sequence (red) to generate a premature stop codon (asterisk) and lead to a truncated protein. Traces represent pooled allele sequences for each individual clone. Two amino acid alterations were detected (S or F). **C**. Immunoblot detection of Gravin (top) and GAPDH (bottom) in wildtype and Gravin KO clonal U2OS cells. **D-F**. Structured illumination microscopy (SIM) images of wildtype (**D**) and Gravin KO (**E & F**) cells during interphase. Composite images (left) show α-tubulin (green) and DAPI (blue). DAPI stain depicted in grayscale (middle). Magnified insets (right) and yellow arrows highlight micronuclei. **G**. Quantification of amalgamated data representing the percent (%) of cells with micronuclei in wildtype (gray) and Gravin KO (red) U2OS cells. Points depict individual experiments (n); wildtype, n=3, Gravin KO, n=3, **p=0.001; A total of 1500 cells were analyzed over three independent experiments. P values were calculated by unpaired two-tailed Student’s t-test. Data are mean ± s.e.m. **H**. Immunoblot detection of Gravin (blot 1), Plk1 (blot 2), γ-tubulin (blot 3), and GAPDH (blot 4) in wildtype and Gravin KO U2OS cells.

We validated our clonal cell lines in five stages. First, Sanger sequencing was used to confirm introduction of indels in Gravin KO clones (Figure 2B). Sequence analysis indicates introduction of a premature stop codon that abolishes the Gravin coding region at target site 1 in our Gravin KO clone (Figure 2B, red sequence). Second, immunoblots detected robust Gravin expression in wildtype cells while a complete loss of the protein was observed in KO lines (Figure 2C). Immunoblot detection of GAPDH served as a loading control (Figure 2C). Removal of Gravin was validated with two separate antibodies against this anchoring protein (Figure 2C and Figure S2A). Third, we assessed our genome-edited clones for the presence of micronucleated cells. Micronuclei are structures that contain genetic material outside of the main nucleus and often result from missegregated chromosomes (28). Our experiments in Gravin-depleted HEK 293 cells demonstrate that micronuclei formation can be used as a functional readout for Gravin loss (Figure S2B). Indeed, we observed this phenomenon in our genetically-engineered human U2OS clones as well (Figure 2D-G). Interphase wildtype cells display a low incidence of micronuclei at baseline (Figure 2D). On the contrary, we observe more micronucleated cells when Gravin is ablated (Figure 2E & F and Figure S2C). Gravin loss results in a 2.4-fold increase in the number of micronucleated cells (Figure 2G). This enrichment was further validated in an additional Gravin KO clone (Figure S2C). Fourth, immunofluorescent detection and quantitative analyses confirmed that levels of active Plk1 (assessed by pT210-Plk1 immunofluorescence signal), a previously-characterized Gravin interacting partner, were reduced at mitotic spindle poles in Gravin KO cells as compared to wildtype controls (Figure S2D-F). Finally, additional immunoblotting established that both wildtype and Gravin KO U2OS cell lines express normal levels of γ-tubulin and Plk1 (Figure 2H). Together, these data provide evidence for an engineered human cell line that lacks the Gravin anchoring protein.

### Gravin deletion enhances the asymmetric distribution of γ-tubulin

Gravin governs the asymmetric distribution of protein kinases at mitotic spindle poles (11). We employed super-resolution structured illumination microscopy (SIM) to examine γ-tubulin organization at each spindle pole. Accordingly, we found that this key microtubule assembly component distributes asymmetrically between the two poles in wildtype MEFs (Figure 3A). This result is further illustrated by representative line plots depicting maximum intensity signals across the x-plane (Figure 3B & C). Quantitative analysis using integrated intensity measurements establish that pole 1 clusters 19.9% more γ-tubulin than pole 2 in wildtype MEF cells (Figure 3D). Parallel analyses recapitulate this phenomenon in wildtype U2OS cells (Figure 3E and Figure S3A). These additional studies demonstrated an 18.9% enrichment of γ-tubulin at pole 1 as compared to pole 2 (Figure 3E). These data reveal the asymmetric distribution of γ-tubulin between individual spindle poles in both wildtype murine and human cells.

**Figure 3:**
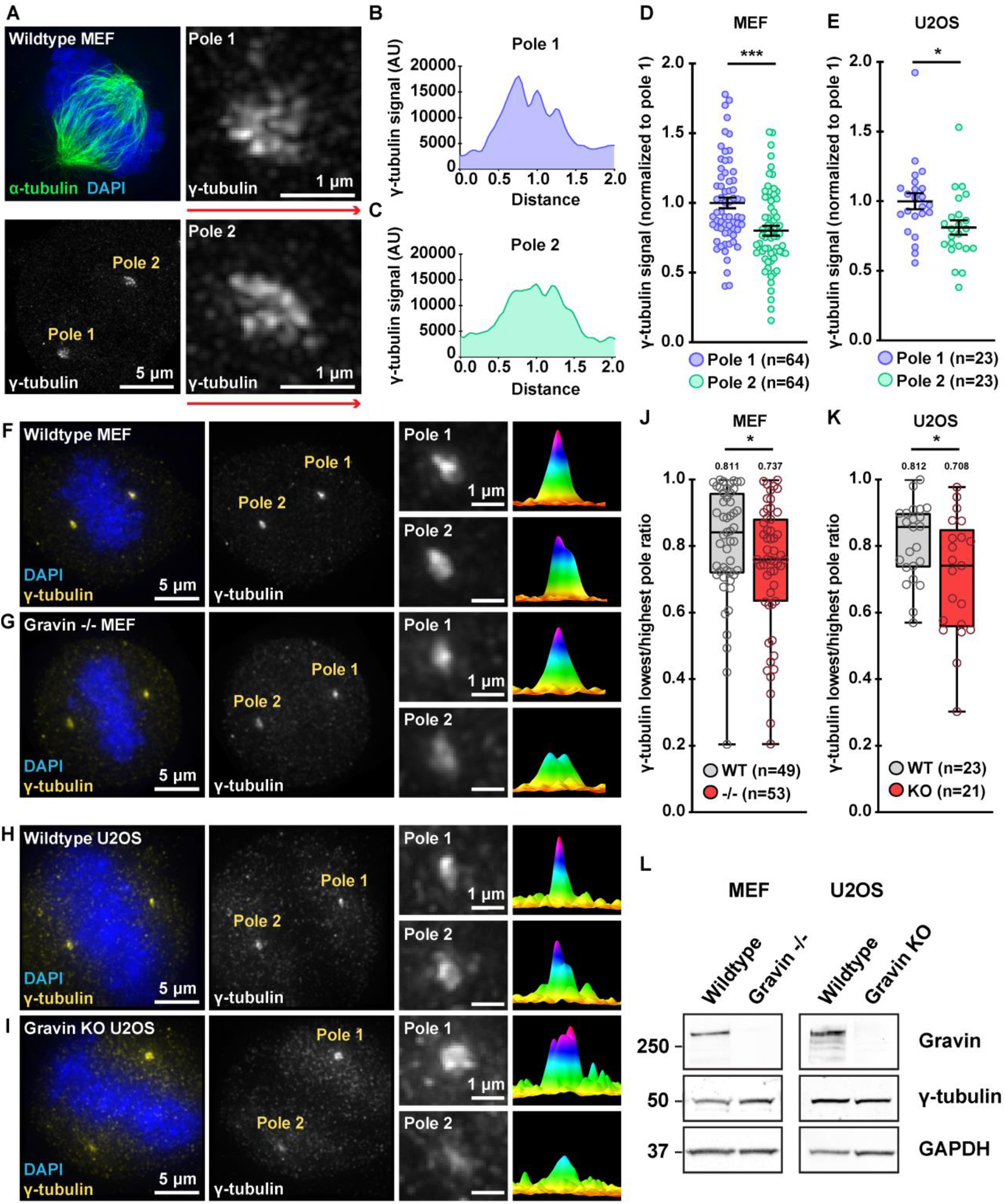
Deletion of Gravin enhances the asymmetric distribution of γ-tubulin. **A**. SIM micrograph of a wildtype MEF during mitosis. Composite images (top left) show α-tubulin (green) and DAPI (blue). Grayscale images depict γ-tubulin at both pole (bottom left). Magnified images reveal that pole 1 (top right) accumulates more γ-tubulin than pole 2 (bottom right). **B & C**. Signal intensity (y-axis) is graphed against distance (x-axis) to generate plot profiles of maximum intensity signal measurements. Distance along red lines in A were used to generate representative plot profiles for pole 1 (**B**) and pole 2 (**C**). **D & E**. Quantification of γ-tubulin immunofluorescence at pole 1 (blue) and pole 2 (green) for MEF (**D**) and U2OS (**E**) mitotic cells; (**D**) pole 1, n=64, pole 2, n=64, ***p=0.0002; (**E**) pole 1, n=23, pole 2, n=23, *p=0.0181. **F-I**. Immunofluorescence of representative wildtype MEF (**F**), Gravin -/-MEF (**G**) wildtype U2OS (**H**) and Gravin KO (**I**) mitotic cells. Composite images (far left) show DAPI (blue) and γ-tubulin (yellow). Grayscale images (middle left) depict γ-tubulin at poles. Magnified insets (middle right) show signals at individual poles. Surface plots (far right) depict signal intensity profiles of each pole. **J & K**. Quantification of γ-tubulin immunofluorescence at each pole represented as box plots showing lowest/highest pole ratio in wildtype (gray) and Gravin null (red) MEF (**J**) and U2OS (**K**) mitotic cells. Mean values are indicated above each plot; (**J**) WT, n=49, -/-, n=53, *p=0.0438; (**K**) WT, n=23, KO, n=21, *p=0.0263. Points in **D, E, J** and **K** represent individual cells (n). Data in **D** and **E** are normalized to pole 1. Experiments were conducted at least three times (N=3). P values were calculated by unpaired two-tailed Student’s t-test. Data are mean ± s.e.m. **L**. Immunoblot detection of Gravin (top), γ-tubulin (middle), and GAPDH (bottom) in wildtype and Gravin null MEF (left) and U2OS (right) cells.

Loss of Gravin disrupts the recruitment of signaling elements to mitotic spindle poles (11,26). Therefore, we tested whether Gravin ablation perturbs the distribution of γ-tubulin between the two poles (Figure 3F-K). As before, immunofluorescent staining in wildtype MEFs confirmed an enrichment of γ-tubulin (yellow) at one spindle pole over the other (Figure 3F). Magnified insets and representative surface plots emphasize this result (Figure 3F). Surprisingly, Gravin null MEFs displayed a more pronounced asymmetry in the pole-to-pole distribution of γ-tubulin (Figure 3G). Quantitative methods were employed to further probe this observation. First, integrated intensity measurements were used to quantitate the total γ-tubulin signal at each pole. Then, the γ-tubulin signal at the pole with the lowest intensity was divided by the signal at the pole with the highest intensity giving us the ratio of γ-tubulin between the poles (“lowest/highest pole ratio”). In wildtype MEFs, we calculated a mean lowest/highest pole ratio of 0.811 (Figure 3J). However, in Gravin null cells we observed an average ratio of 0.737 (Figure 3J). Parallel analyses in the CRISPR/Cas9-edited U2OS cells revealed a similar result (Figure 3H, I & K). Wildtype U2OS cells displayed a minor enrichment of γ-tubulin at one pole over the other (Figure 3H). Quantitative analysis of multiple wildtype cells yielded a mean lowest/highest pole ratio of 0.812 (Figure 3K). Consistent with our earlier findings, Gravin KO U2OS cells displayed a ratio of 0.708. This asymmetry was further validated in an additional Gravin KO clone (Figure S3B). Finally, immunoblots established that total γ-tubulin expression was similar between wildtype and Gravin null genotypes in both MEF and U2OS cell lines (Figure 3L). Collectively, these findings suggest that Gravin deletion leads to enhanced asymmetry of γ-tubulin between mitotic spindle poles.

### Precision targeting of Plk1 inhibitors to spindle poles promotes asymmetric distribution of active Plk1 and γ-tubulin

Gravin anchors Plk1 during mitosis and depletion of this scaffold reduces the pool of active kinase at the mitotic spindle poles (11,26). Thus, it is possible that reduced Plk1 activity may contribute to the defects seen in our Gravin null U2OS cells. We previously reported that treatment with the Plk1 inhibitor BI2536 reduces the total levels of pT210-Plk1 (an index of active kinase) at centrosomes (26). Here we examined how the pT210-Plk1 pool that remains after inhibition with BI2536 distributes between the individual poles. In DMSO-treated controls, immunofluorescent detection of pT210-Plk1 revealed a mild enrichment of active kinase at one spindle pole over the other (Figure 4A). Remarkably, inhibition of Plk1 led to greater asymmetry in the pole-to-pole distribution of pT210-Plk1 (Figure 4A). This effect is even more pronounced with increasing concentrations of Plk1 inhibitor (Figure 4A). Representative line plots further illustrate this result (Figure 4B). Integrated intensity measurements were used to quantitate the total pT210-Plk1 signal at the spindle poles for each inhibitor dose (Figure 4C). While DMSO-treated cells display an 18.7% enrichment in pT210-Plk1 at one pole over the other, cells treated with 20 nM BI2536 show a 67.8% difference between the two poles (Figure 4C). These data suggest that inhibition of Plk1 perturbs the distribution of active kinase between the spindle poles.

**Figure 4:**
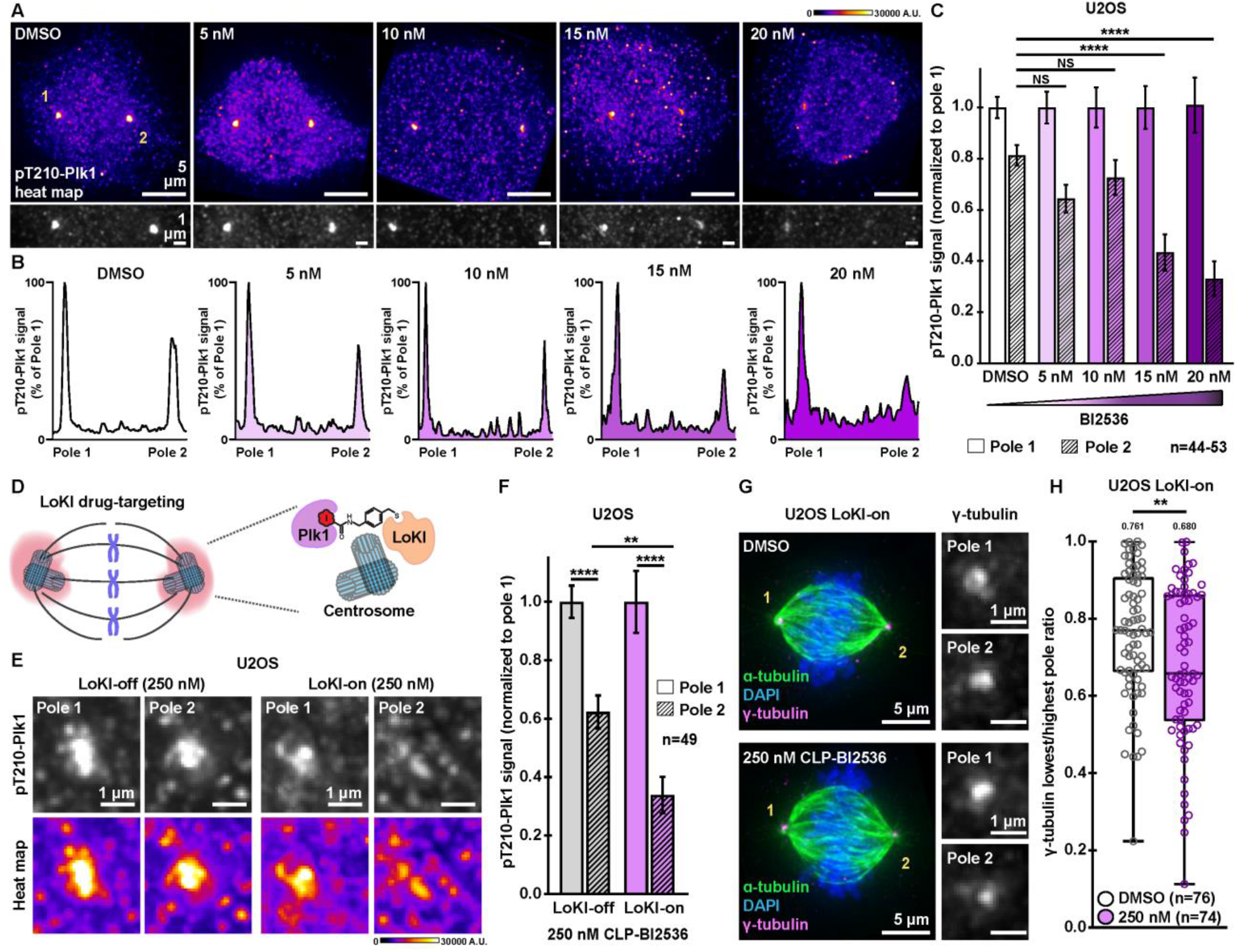
Targeting Plk1 inhibitors to spindle poles promotes a more asymmetric distribution of active Plk1 and γ-tubulin. **A**. Immunofluorescence detection of pT210-Plk1 (an index of kinase activity) at mitotic spindle pole in parental U2OS cells treated with DMSO or BI2536 for 4 hr. Distribution of pT210-Plk1 is represented with pseudo-color heat maps (top) and in grayscale (bottom). Signal intensity scale (A.U.) is shown above. **B**. pT210-Plk1 signal (grayscale from A) is graphed against distance to generate plot profiles of maximum intensity signal measurements. Peaks are normalized to pole 1. BI2536 concentrations are indicated. **C**. Quantification of amalgamated pT210-Plk1 immunofluorescence at each pole for U2OS parental cells treated with DMSO or BI2536; pole 2, DMSO, n=53, control; 5nM, n=44, p=0.1387; 10 nM, n=44, p=0.7016; 15 nM, n=44, ****p<0.0001; 20 nM, n=44, ****p<0.0001. **D**. Schematic of precision drug targeting of Plk1 inhibitor to spindle poles (pink) using the Local Kinase Inhibition (LoKI) system. **E**. Immunofluorescence of pT210-Plk1 at individual spindle poles in representative LoKI-off (left) and LoKI-on (right) U2OS cells treated with 250 nM CLP-BI2536 for 4 hr. Signal of active kinase is represented in grayscale (top) and with pseudo-color heat maps (bottom). Signal intensity scale (A.U.) is shown below. **F**. Quantification of amalgamated pT210-Plk1 immunofluorescence at each pole after 4 hr of targeted BI2536 delivery; LoKI-off, pole 1, n=49, LoKI-off, pole 2, n=49, ****p<0.0001; LoKI-on, pole 1, n=49, LoKI-on, pole 2, n=49, ****p<0.0001; LoKI-off, pole 2, n=49, LoKI-on, pole 2, n=49, **p=0.0010. **G**. Immunofluorescence of a LoKI-on mitotic cell treated with DMSO (top) or 250 nM CLP-BI2536 (bottom) for 4 hr. Composite images (left) show α-tubulin (green), DAPI (blue), and γ-tubulin (magenta). Magnified grayscale images (right) show γ-tubulin signal at poles. **H**. Quantification of amalgamated γ-tubulin immunofluorescence data at each pole. Data is represented as box plots showing lowest/highest pole ratio for LoKI-on cells after 4 hr treatment with DMSO (white) or 250 nM CLP-BI2536 (purple) followed by a 1 hr washout. Mean values are indicated above each plot; DMSO, n=76, 250 nM, n=74, **p=0.0074. Values depicted in **C**, **F** and **H** represent individual cells (n). Data in **C** and **F** are normalized to pole 1. Experiments were conducted at least three times (N=3). P values were calculated by one-way ANOVA with Dunnett’s multiple comparisons test performed with pole 2 DMSO as control (**C**) or unpaired two-tailed Student’s t-test (**F** and **H**). Data are mean ± s.e.m. NS, not significant.

Since Gravin anchors Plk1 at mitotic spindle poles we assessed whether selective inhibition of Plk1 at this location disrupts the pole-to-pole distribution of active kinase. This was achieved by Local Kinase Inhibition (LoKI), a drug-targeting approach that allows us to deliver the Plk1 inhibitor CLP-BI2536 to centrosomes (26). This precision pharmacology tool allowed us to inhibit Plk1 specifically at spindle poles (Figure 4D). Immunofluorescence detection of pT210-Plk1 assessed the levels of active kinase at each spindle pole (Figure 4E and Figure S4A). Control cells express a mutant construct called LoKI-off that is unable to conjugate derivatized kinase inhibitors at spindle poles. When treated with 250 nM CLP-BI2536, these cells display 37.7% less pT210-Plk1 (gray) at pole 2 as compared to pole 1 (Figure 4E & F). Heat plots further emphasize this result (Figure 4E). Strikingly, cells expressing the active LoKI-on targeting platform exhibited 66.1% less pT210-Plk1 at pole 2 as compared to pole 1 when treated with the same concentration of drug (Figure 4E & F). Quantitative analyses established a 1.8-fold greater difference in the distribution of active kinase with spindle pole-targeted inhibitors as comparted to global drug (Figure 4F). Additional controls were conducted in DMSO-treated U2OS cells (Figure S4A & B). These studies confirmed that in the absence of drug we observe comparable distributions of pT210-Plk1 between the poles in cells expressing either LoKI targeting platform (Figure S4A & B). Additional validation confirmed that the enhanced asymmetry of pT210-Plk1 does not result from redistribution of total Plk1 protein (Figure S4C). Together, these data suggest that local targeting of Plk1 inhibitor drugs to mitotic spindle poles leads to enhanced asymmetry in the distribution of active kinase at the spindle poles.

Recruitment of γ-tubulin to centrosomes relies on Plk1 activity (29,30). Consequently, inhibition of Plk1 at this location disrupts γ-tubulin targeting to these structures (26). Thus, a logical next step was to test whether local inhibition of Plk1 enhances the asymmetrical localization of γ-tubulin between the individual spindle poles. Targeted delivery of CLP-BI2536 in LoKI-on cells promoted a more asymmetric distribution of γ-tubulin as compared to DMSO controls (Figure 4G). Magnified insets of each pole further emphasize this result (Figure 4G). Quantitative analysis revealed that the lowest/highest pole ratio for γ-tubulin drops from 0.761 to 0.680 when Plk1 inhibitors are targeted to the mitotic spindle poles (Figure 4H). In contrast, a less pronounced effect was observed when experiments were repeated in LoKI-off cells (Figure S4D & E). These precision pharmacology experiments reveal that targeted Plk1 inhibitor drugs lead to an enhanced asymmetric localization of γ-tubulin at mitotic spindle poles.

### Gravin ablation diminishes interactions between γ-tubulin and upstream regulators

Nedd1, a member of the γ-tubulin ring complex (γ-TURC), coordinates γ-tubulin accumulation at centrosomes (31,32). Furthermore, Plk1 promotes the interaction of γ-tubulin with Nedd1 to ensure proper targeting of γ-TURC during mitosis (33). We employed proximity ligation assay (PLA), a technique that marks protein-protein interactions that occur within a range of 40–60 nm, to identify protein interaction pairs in wildtype and Gravin null mitotic cells (Figure 5A-E). Immunofluorescent detection of PLA puncta (green) and DAPI (blue) uncovered Plk1/γ-tubulin interaction pairs in wildtype MEFs (Figure 5A). Significantly fewer PLA puncta were detected in Gravin null cells (Figure 5B). Quantitative analysis reveals a 24% reduction in the number of puncta observed in Gravin null MEFs as compared to wildtype controls (9.8 versus 12.9, respectively; Figure 5C). Complementary experiments were carried out to assess Nedd1/γ-tubulin protein-protein interactions (Figure 5D-E). Again, immunofluorescent detection identified PLA puncta (yellow) and DAPI (blue) in mitotic cells (Figure 5D). As before, Gravin null MEFs display fewer PLA puncta than wildtype control cells (Figure 5D). Quantitation further revealed that Gravin-depleted cells display 34.8% less PLA puncta than wildtype MEFs (Figure 5E). These findings suggest that Plk1/γ-tubulin and Nedd1/γ-tubulin interactions are reduced in cells lacking Gravin. Thus, abrogation of Plk1-mediated signaling may underlie the abnormalities in γ-tubulin accumulation observed in Gravin null cells (Figure 5F).

**Figure 5:**
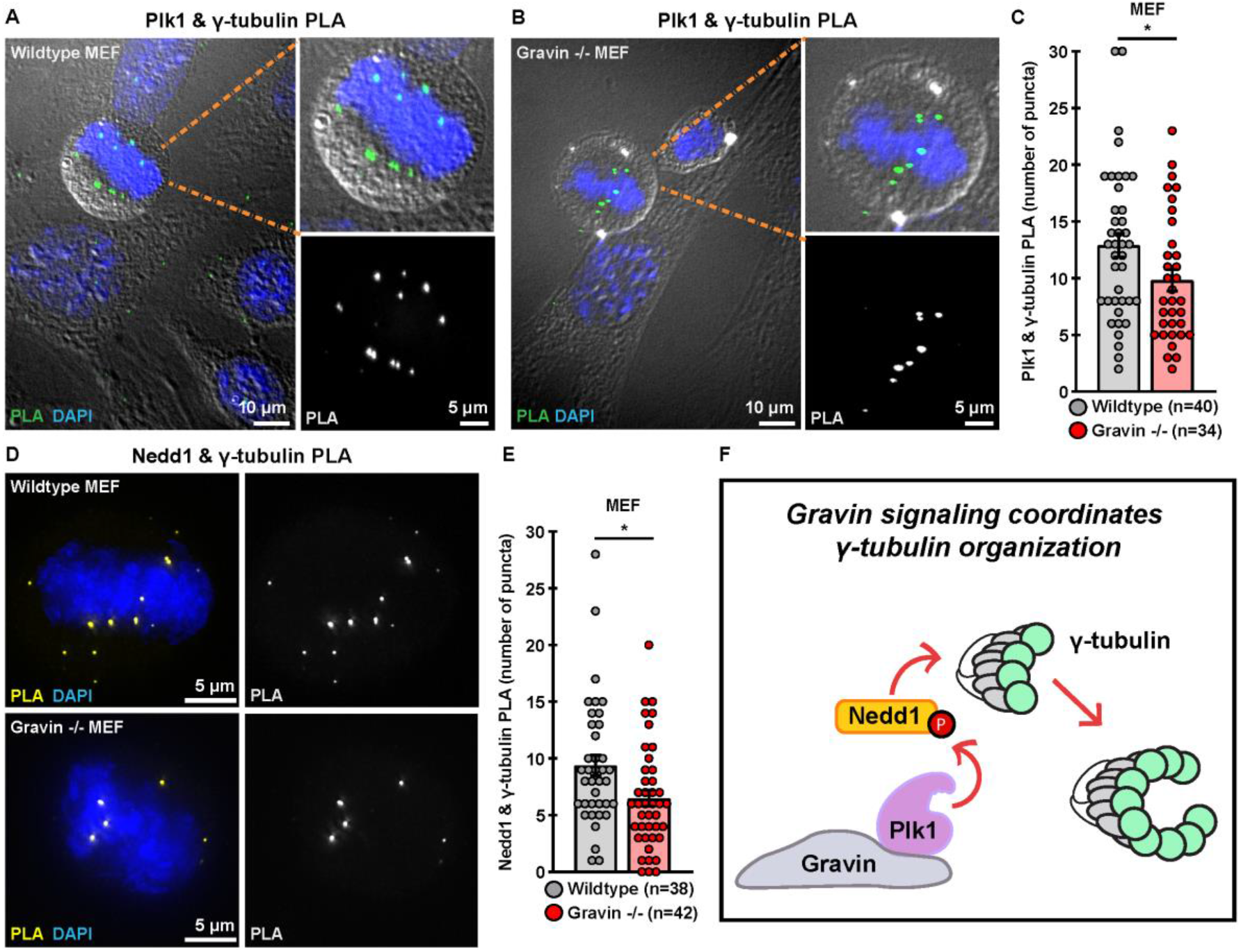
Interactions between γ-tubulin and upstream regulators are disrupted in Gravin-ablated cells. **A & B**. Wildtype (**A**) and Gravin -/- (**B**) MEFs assayed with proximity ligation assay (PLA) reveal Plk1 & γ-tubulin interaction pairs. Composite images (left) show PLA puncta (green) and DAPI (blue). Differential interference contrast (DIC) shows cell boundaries in mitotic (foreground) and interphase (background) cells. Magnified insets of composite images (top right) depict a mitotic cell. Grayscale images (bottom right) reveal PLA puncta. **C**. Quantification of Plk1 & γ-tubulin PLA (number of puncta per cell) in wildtype (gray) and Gravin -/-(red) MEFs; wildtype, n=40, Gravin -/-, n=34, *p=0.0357. **D**. Immunofluorescence of wildtype (top) and Gravin -/-(bottom) MEFs assayed with PLA reveal Nedd1 & γ-tubulin interaction pairs. Composite images (left) show PLA (yellow) and DAPI (blue). Single channel images of PLA are represented in grayscale (right). **E**. Quantification of Nedd1 & γ-tubulin PLA (number of puncta per cell) in wildtype (gray) and Gravin -/-(red) MEFs; wildtype, n=38, Gravin -/-, n=42, *p=0.0131. Points in **C** and **E** represent individual cells (n). Experiments were conducted three times (N=3). P values were calculated by unpaired two-tailed Student’s t-test. Data are mean ± s.e.m. **F**. Schematic depicting that Gravin organizes a signaling network which promotes accumulation of γ-tubulin (gray circles) when Plk1 and Nedd1 are in proximity to γ-tubulin.

## Discussion

Subcellular targeting and anchoring of protein kinases is a recognized molecular mechanism that enhances the precision and fidelity of protein phosphorylation events (3). Gravin regulates various aspects of mitosis including the clustering of mitotic protein kinases to facilitate spindle formation and mitotic progression (11,34,35). In this study, we discover a new role for this anchoring protein in modulating the recruitment of γ-tubulin, a major regulator of microtubule nucleation. Our imaging studies in Figure 1 reveal that loss of Gravin perturbs the accumulation of γ-tubulin at mitotic spindle poles. Elevated expression and ectopic cellular distribution of γ-tubulin has been observed in astrocytic gliomas (17). Delocalization of this key microtubule-nucleating component has also been detected in human breast cancer, with the most dramatic changes occurring in tumor cell lines that encompass the greatest metastatic potential (19). Both reports conclude that inappropriate subcellular distribution of γ-tubulin is associated with tumor progression. Our findings argue that Gravin-mediated signaling further contributes to this process. Other investigators have shown that Gravin loss is linked to defective mitoses in a variety of cancer cell lines (35,36). Recent phosphoproteomics studies further implicate this anchoring protein as a potential prognostic biomarker for high-grade meningiomas (35,36). Our discovery that Gravin-ablated cells display an enhanced pole-to-pole asymmetry of γ-tubulin provides additional clues into how protein scaffolds safeguard cell division events (Figure 3F-K). Thus, we postulate that Gravin may serve a protective role during mitosis by ensuring the proper localization of γ-tubulin at mitotic spindle poles. Collectively, these studies suggest that precise targeting of mitotic signaling components contributes to the fidelity of cell division.

Polo-like kinase 1 is a promising target for therapeutic intervention in cancer (37–41). This enzyme catalyzes phosphorylation events necessary for the efficient and accurate progression of cells through mitosis (42–44). While ATP-competitive kinase inhibitors such as BI2536 are putative anticancer compounds that have entered clinical trials, conventional approaches for delivering these drugs increase off-target effects and mask contribution of Plk1 at specified subcellular locations (37,45,46). With this in mind we reasoned that manipulating molecular scaffolds such as Gravin could offer new insight into how locally-constrained signaling enzymes are regulated. Moreover, pharmacological strategies that inhibit kinase activity within a specified locale permit the investigation of individual signaling events. Here, we demonstrate an application of the new precision pharmacology tool, LoKI, by targeting kinase inhibitor drugs to the subcellular location where Gravin anchors Plk1 (26,47). The rationale for utilizing LoKI targeting was provided by previous reports that have implicated a role for Plk1 in organizing γ-tubulin at mitotic centrosomes (26,30,31,48). A key advance, illustrated in Figure 4, is that suppression of Plk1 activity at mitotic spindle poles enhances the asymmetric distribution of γ-tubulin. By taking advantage of the spatial resolution afforded by the LoKI system we discover that selective loss of Plk1 action at spindle poles perturbs the distribution of γ-tubulin. In addition, the data in Figure 4E-H uncover a link between loss of Plk1 activity at the spindle poles and an enhanced asymmetric distribution of γ-tubulin. On the basis of these results, it is possible that dampened Plk1 activity at mitotic spindle poles underlies the γ-tubulin accumulation defects observed in cells lacking Gravin. Alternatively, Gravin may facilitate phosphorylation of yet unknown mitotic substrates that regulate γ-tubulin targeting. Irrespective of either mechanism, our findings advance the concept that Gravin-anchored kinase activity is necessary to optimally regulate the distribution of a central γ-tubulin ring complex (γ-TURC) component during mitosis.

AKAPs constrain enzymes within spatially restricted signaling islands to restrict kinases and phosphatases to the immediate vicinity of select substrates (49–51). Our previous reports showed that Gravin organizes Plk1 and Aurora A kinases at spindle poles to facilitate proper spatial relay of signaling during mitosis (11,26). A new concept emerging from this study is that cells without Gravin have lost the necessary protein-protein interactions to drive robust localization of γ-tubulin at spindle poles. This is supported by data in Figure 5D-E showing that γ-tubulin interaction with Nedd1, a Plk1 substrate, is disrupted in Gravin null cells. On the basis of these findings we conclude that inappropriate targeting of Plk1 activity in Gravin-ablated cells may contribute to the loss of γ-tubulin/Nedd1 interactions. Moreover, our data is in line with a recent study which discovered that increased Plk1 mobility in Gravin-depleted cells results in aberrant phosphorylation of CEP215, a PCM component that regulates the assembly of the γ-tubulin ring complex (γ-TURC) (25). Since Plk1 promotes the interaction of γ-tubulin with Nedd1, it is possible that loss of this interaction in Gravin null cells drives the disruptions in γ-tubulin accumulation that we observe (33). Thus, our findings in Figure 5 offer additional mechanistic insight into how Gravin organizes signaling complexes at specific subcellular location to ensure that kinases are positioned in close proximity to their targets.

A recognized facet of AKAPs is their ability to work within spatially restricted microenvironments to direct and insulate kinase action within a few angstroms of their intended targets (49,52). Kinase anchoring is particularly effective for the modulation of a highly coordinated process such as the cell cycle (53). Thus, it is perhaps not surprising that subtle perturbations in the organization of key macromolecular complexes are responsible for certain defective signaling events that are observed in a range of cancers (Bucko and Scott, in press). This is evidenced by studies that have identified the *AKAP12* gene, which encodes Gravin, in a deletion hotspot for a variety of human cancers (54). However, it is equally worth noting that elevated levels of Gravin in ovarian cancer have been correlated with poor prognosis (55). Thus, more studies are necessary to clarify the role of Gravin in these different pathological contexts. Another theme emerging from our work is that deploying tools that direct kinase inhibitors to specific subcellular locations provide clues into the contributions of local signaling events. This study provides a practical application of the LoKI system and demonstrates that precision pharmacology can be utilized to decipher how anchored Plk1 activity at the spindle poles drives the asymmetric distribution of γ-tubulin. Together, our findings lead us to speculate that perturbation of Gravin signaling during mitosis may underlie disease progression. However, future work is necessary to uncover additional Gravin binding partners and downstream substrates that become dysregulated when this anchoring protein is ablated in disease contexts.

## Experimental procedures

### Antibodies and reagents

Alpha-tubulin, clone DM1A (Sigma Aldrich, T9026, mouse monoclonal; RRID: AB_477593); Alpha-tubulin-FITC, clone DM1A (Sigma Aldrich, F2168, mouse monoclonal; RRID: AB_477593); Flag M2 (Sigma Aldrich, F3165, mouse monoclonal, RRID: AB_259529); Flag-HRP (Sigma Aldrich, A8592, mouse monoclonal RRID: AB_439702); Gamma-tubulin (Abcam, 11317, rabbit polyclonal; RRID: AB_297921); GAPDH-HRP (Novus, NB110-40405, mouse monoclonal; RRID: AB_669249); Gravin, clone JP74 (Sigma Aldrich, G3795, mouse monoclonal); Gravin, clone R3698 (in-house (34), rabbit polyclonal); Nedd1, clone H-3 (Santa Cruz Biotechnology, sc-398733, mouse monoclonal); Pericentrin (Abcam, ab4448, rabbit polyclonal; RRID:AB_304461); Phospho-Plk1 T210 (Biolegend, 628901, rabbit polycolonal; RRID: AB_439786); Phospho-Plk1 T210 clone 2A3 (Abcam, ab39068, mouse monoclonal, RRID: AB_10861033); Plk1, clone 35-206 (Millipore, 05-844, mouse monoclonal; RRID: AB_11213632); Plk1, clone F-8 (Santa Cruz Biotechnology, sc-17783, mouse monoclonal; RRID: AB_628157); Ssecks (BD Transduction Labs, S63820, mouse monoclonal). Donkey anti-Mouse IgG, Alexa Fluor 488 (Invitrogen, A-21202); Donkey anti-Rabbit IgG, Alexa Fluor 647 (Invitrogen, A-31573). Amersham ECL Mouse IgG, HRP-linked F(ab’)₂ fragment from sheep (GE Life Sciences, NA9310); Amersham ECL Rabbit IgG, HRP-linked F(ab’)₂ fragment from donkey (GE Life Sciences, NA9340).

### Plasmid constructs

Constructs for the generation of LoKI cells were generated as described in (26). In brief, LoKI constructs contain an N-terminal mCherry reporter protein, two SNAP-tag domains, and a PACT (AKAP450 centrosomal targeting) sequence. The second SNAP-tag has a varied codon sequence to simplify PCR-amplification. Individual components were PCR amplified with overlapping ends and/or Gateway “att” sites and assembled using Gibson Cloning. Constructs were subcloned into pLIX402 (a gift from David Root; Addgene plasmid #41394) using Gateway Cloning. Mutant LoKI vectors were generated by performing site-directed mutagenesis with a QuikChange II XL kit (Aligent). Constructs for CRISPR/Cas9-mediated genome editing experiments expressing Cas9 and gRNAs were generated by cloning into pSpCas9(BB)-2A-Puro (PX459) vectors (Addgene plasmid #48139). Constructs were verified by Sanger sequencing.

### Cell culture

U2OS cells used to generate stable cell lines (26) were purchased from ATCC and tested negative for mycoplasma contamination as assessed by the Universal Mycoplasma Detection Kit (ATCC 30-1012K). U2OS, Control and Gravin shRNA HeLa (11), Control and Gravin shRNA HEK293 (10) and immortalized MEF (generated as described in (11)) cells were maintained in DMEM, high glucose (Life Technologies) at 37°C and 5% CO2. All media was supplemented with 10% FBS (Thermo Fisher). Infections for generation of stable knockdown in HeLa and HEK293 cells were performed with shRNA lentiviral particles (Santa Cruz Biotech). Infections for generation of stable LoKI cells were performed using lentiviral particles created in-house (as described in (26)). Transient gene expression of Flag-Ssecks, a murine Gravin rescue construct, was performed by transfection using TransIT-LTI reagent (Mirus).

### CRISPR-Cas9 editing of Gravin

CRISPR/Cas9-mediated genome editing was used to delete the AKAP12 gene on chromosome 6 in U2OS cells. Two unique guide RNAs (gRNAs) were designed to the following targets: gRNA1, AGAGATGGCTACTAAGTCAG**CGG**; gRNA 2, AGCCGAATCTGGCCAAGCAG**TGG**. Bold letters indicate PAM sequence. Constructs expressing Cas9 and either gRNA were generated by cloning into pSpCas9(BB)-2A-Puro (PX459) vectors (Addgene plasmid #48139). Individual vectors were transfected into U2OS cells using TransIT-LT1 reagent (Mirus) in Opti-MEM® (Life Technologies) media according to manufacturer’s instructions. Single clones were isolated using Scienceware cloning discs (Sigma Aldrich). Clones were first screened by immunoblotting for loss of Gravin protein expression using two independent antibodies. Clones that had undetectable levels of Gravin were further checked for mutations. In brief, cells were pelleted and purification of genomic DNA was achieved by treating with 50 mM NaOH for 20 minutes at 95°C followed by treatment with 1M Tris pH 8.0 and a 10 minutes spin at 14,000 rpm. Primers flanking the target sites were designed and the region was amplified by PCR: gRNA1-forward, TTGGACAGAGAGACTCTGAAGATGTG; gRNA1-reverse, CTTCATCTTTCTTCACAGTGAGTAGC; gRNA2-forward, ACTGTGAAGAAAGATGAAGGGG. PCR products were cloned into vectors using the Zero Blunt™ TOPO™ PCR Cloning Kit (Thermo Fisher). Individual clones were isolated for plasmid DNA mini-prep and verified by Sanger sequencing.

### Drug treatments

RO3306 (Sigma) was used to enrich for mitotic cells in Gravin rescue experiments. At least 24 hours post transfection, cells were treated with 10 μM RO3306 for 24 hours and then incubated in complete DMEM for 25 minutes prior to fixation. BI2536 (AdooQ) and CLP-BI2536 (26) were used to inhibit Plk1. For LoKI experiments, cells were treated as described in (26). In brief, prior to drug treatment, cells were incubated with 1 μg/mL doxycycline hyclate (Sigma-Aldrich,) in FBS-supplemented DMEM for 48-72 hours to induce expression of LoKI targeting platforms. Cells were grown on 1.5 poly-D-lysine coated coverslips (neuVitro) for ~16 hours in complete DMEM and then treated with Dimethylsulfoxide, DMSO (Pierce) or CLP-BI2536 in serum-free DMEM for 4 hours. For washout experiments (γ-tubulin data), cells were incubated in serum-free DMEM without inhibitors for an additional 1 hour. Cells were washed once with PBS prior to fixation.

### Immunoblotting

Cells were lysed in RIPA buffer (50 mM Tris HCl pH 7.4, 1% Triton X-100, 0.5% Sodium Deoxycholate, 0.1% SDS, 50 mM NaF, 120 mM NaCl, 5 mM β-glycerophosphate) with protease and phosphatase inhibitors (1 mM benzamidine, 1 mM AEBSF, 2 μg/mL leupeptin, 100 nM microcystin-LR). Samples were boiled for 5 minutes at 95°C in NuPAGE™ LDS Sample Buffer 4X (Thermo Fisher) + 5% BME (Sigma-Aldrich) and protein concentration was determined using a Pierce™ BCA Protein Assay Kit (Thermo Fisher). Samples were resolved on Bolt® 4-12% Bis-Tris Plus Gels (Invitrogen), proteins were transferred to nitrocellulose for immunoblotting, and membranes were probed with primary antibodies. Detection was achieved with a HRP-conjugated rabbit or mouse secondary antibody (GE Healthcare) followed by enhanced chemiluminescence with SuperSignal™ West Pico PLUS Chemiluminescent Substrate (Thermo Fisher). Representative blots were adjusted for brightness and contrast in Fiji.

### Immunofluorescence

Cells grown on 1.5 poly-D-lysine coated coverslips (neuVitro) for ~16 hours were fixed in ice-cold methanol or in 4% paraformaldehyde in PBS for 10 minutes. Cells were permeabilized and blocked in PBS with 0.5% Triton X-100 and 1% BSA (PBSAT) for 30 minutes. Primary antibodies and secondary antibodies, conjugated to Alexa Fluor dyes (Invitrogen), were diluted in PBSAT and cells were stained for 1 hour in each. Counterstaining with FITC-tubulin antibodies and/or DAPI (Thermo Fisher) was carried out for 10-45 minutes in PBSAT. Washes with PBSAT were carried out in one of two ways (3X for 5 minutes or 10X, quick on and off) between antibody and/or dye incubation steps and prior to mounting. Coverslips were mounted on slides using ProLong® Diamond Antifade Mountant (Life Technologies, P36961).

### Proximity Ligation Assay

Cells were methanol-fixed, permeabilized, blocked, and stained with primary antibodies as described under “immunofluorescence”. Cells were incubated with anti-rabbit and anti-mouse probes (Duolink) and PLA was carried out according to manufacturer’s instructions. Where applicable, cells were counterstained as described above.

### Microscopy

Widefield and super-resolution 3D-SIM images were acquired on a Deltavision OMX V4 (GE Healthcare) system equipped with a 60x/1.42 NA PlanApo oil immersion lens (Olympus), 405-, 488-, 568-, and 642-nm solid-state lasers and sCMOS cameras (pco.edge). For SIM, 15 images per optical slice (3 angles and 5 phases) were acquired. Image stacks of 4-7 μm with 0.200 (widefield) or 0.125-μm (SIM) optical thick z-sections were acquired using immersion oil with a refractive index 1.516 or 1.518. Z-stacks were generated using the DAPI or α-tubulin channels to define the upper and lower boundaries of the plane with a 0.5 μm step size. SIM images were reconstructed using Wiener filter settings of 0.003 and optical transfer functions measured specifically for each channel with SoftWoRx software (GE Healthcare) were used to obtain super-resolution images with a two-fold increase in resolution both axially and laterally. Images from different color channels were registered using parameters generated from a gold grid registration slide (GE Healthcare) and SoftWoRx. Widefield images were deconvolved using SoftWoRx. For micronuclei counts and acquisition of DIC images for PLA assays, a DM16000B inverted microscope (Leica) equipped with a 63x Plan-Apocromat NA 1.4 oil objective, a CSU10 confocal spinning disk (Yokogawa) and a CoolSnap HQ camera (Photometrics) controlled by MetaMorph 7.6.4 (Molecular Devices) was used. Representative images were adjusted for brightness and contrast in Fiji.

### Image Analysis

SoftWoRx (GE Healthcare) or NIH ImageJ (Fiji) software was used to generate maximum intensity projections from z-stack images. Immunofluorescence signals were measured using Fiji software. For analysis of immunofluorescence at spindle poles, sum slice 32-bit Tiff projections were generated from z-stack images. The oval selection tool in Fiji was used to draw a circle (ROI) around the spindle pole and measure the signal in the 647 (γ-tubulin) channel. The area of the circle remained consistent for all experimental replicates. Using the measure function in Fiji, with “Area” and “Raw Integrated Density” predefined as measurements, values were recorded for each spindle pole and for a nearby background region.

Total signal at spindle poles: The average raw integrated density for the spindle poles was determined by adding together the raw integrated densities for each pole in a cell and dividing that value by two. The integrated density for the background was subtracted from the average spindle pole integrated density to yield a background-subtracted average integrated density signal. In cases of a negative value (when background signal is higher than that at poles), a value of zero was reported. An average spindle pole signal was calculated for each control and experimental condition. To do this, normalized average integrated densities were added together and divided by the total number of cells for a given condition. This resulted in a value representing the background-normalized average integrated density at the spindle poles. Values for Gravin null and drug-treated cells were normalized to wildtype or DMSO-treated controls, respectively.

Signal at individual spindle poles: The individual raw integrated densities for each spindle pole in a given cell were classified into two categories: pole 1 (the spindle pole that contains the highest fluorescence signal) and pole 2 (the pole with the lowest fluorescence signal). The background signal values were subtracted from each individual pole. As before, negative values were replaced with a zero. Values for the low pole were normalized to average high pole values of each corresponding condition.

Lowest/highest pole ratios: Background-subtracted integrated intensity values were used to determine the total γ-tubulin signal at each pole. The immunofluorescence signal of the pole with the lowest intensity was divided by the signal of the pole with the highest intensity to determine the ratio of γ-tubulin between the poles. In cases where the value for the ratio was zero, the value was removed and not included in statistical analysis. This was to avoid misinterpretation of the data since a value of zero at both poles and a high value at pole 1 but a zero value at pole 2 would both yield the same ratio. Micronuclei analysis: For each experimental replicate, 500 interphase cells were examined using the DAPI channel. The number of cells with micronuclei was divided by 500 to determine the percent of micronucleated cells per experiment.

PLA analysis: The number of PLA puncta per cell was quantified using the cell counter tool in Fiji.

Surface and line plots: Plots were generated in Fiji software from maximum intensity projections of representative images using the 3D Surface Plot function or the Plot Profile function. The Image J “fire” LUT setting was used to generate pseudo-color images.

### Statistical analysis

Statistics were performed using an unpaired two-tailed Student’s t-test or a one-way ANOVA in Prism 8 software (Graphpad). All values are reported as mean ± standard error of the mean (s.e.m) with p-values less than 0.05 considered statistically significant. Number of independent experiments (N) and number of individual points over several experiments (n) are presented. The sample size was not statistically determined. Where applicable, n > 20 independent measurements were conducted across N ≥ 3 independent experiments. For micronuclei experiments in Gravin KO #2 (Figure S2C) at least 1000 cells per condition over 2 independent experiments were assayed. Detailed analyses are presented in Table 1.

**Table 1:** Statistical Analyses Table.

| Figure | Condition | Sample | Test | Statistics | p-value | Summary |
| --- | --- | --- | --- | --- | --- | --- |
| 1H | Wildtype | 49 | unpaired two-tailed Student's t-test | t=5.855, df=101 | <0.0001 | **** |
|  | Gravin -/- | 54 |  |  |  |  |
| S1C | Control shRNA | 46 | unpaired two-tailed Student's t-test | t=3.901, df=90 | 0.0002 | *** |
|  | Gravin shRNA | 46 |  |  |  |  |
| S1E | Control shRNA | 30 | unpaired two-tailed Student's t-test | t=8.145, df=57 | <0.0001 | **** |
|  | Gravin shRNA | 29 |  |  |  |  |
| S1E | Gravin shRNA | 29 | unpaired two-tailed Student's t-test | t=6.217, df=57 | <0.0001 | **** |
|  | Gravin shRNA + rescue | 30 |  |  |  |  |
| S1E | Control shRNA | 30 | unpaired two-tailed Student's t-test | t=1.821, df=58 | 0.0737 | NS |
|  | Gravin shRNA + rescue | 30 |  |  |  |  |
| 2G | Wildtype | 3 | unpaired two-tailed Student's t-test | t=8.527, df=4 | 0.0010 | ** |
|  | Gravin KO | 3 |  |  |  |  |
| S2B | Control shRNA | 3 | unpaired two-tailed Student's t-test | t=5.340, df=4 | 0.0059 | ** |
|  | Gravin shRNA | 3 |  |  |  |  |
| S2C | Wildtype | 2 | unpaired two-tailed Student's t-test | t=26.90, df=2 | 0.0014 | ** |
|  | Gravin KO | 2 |  |  |  |  |
| S2F | Wildtype | 42 | unpaired two-tailed Student's t-test | t=5.391, df=79 | <0.0001 | **** |
|  | Gravin KO | 39 |  |  |  |  |
| 3D | Pole 1 | 64 | unpaired two-tailed Student's t-test | t=3.794, df=126 | 0.0002 | *** |
|  | Pole2 | 64 |  |  |  |  |
| 3E | Pole 1 | 23 | unpaired two-tailed Student's t-test | t=2.454, df=44 | 0.0181 | * |
|  | Pole2 | 23 |  |  |  |  |
| 3J | WT | 49 | unpaired two-tailed Student's t-test | t=2.042, df=100 | 0.0438 | * |
|  | -/- | 53 |  |  |  |  |
| 3K | WT | 23 | unpaired two-tailed Student's t-test | t=2.303, df=42 | 0.0263 | * |
|  | KO | 21 |  |  |  |  |
| S3B | WT | 23 | unpaired two-tailed Student's t-test | t=2.303, df=42 | 0.0263 | * |
|  | KO | 21 |  |  |  |  |
| S3B | WT | 23 | unpaired two-tailed Student's t-test | t=2.452, df=37 | 0.0190 | * |
|  | KO #2 | 16 |  |  |  |  |
| S3B | KO | 21 | unpaired two-tailed Student's t-test | t=0.2261, df=35 | 0.8225 | NS |
|  | KO #2 | 16 |  |  |  |  |
| 4C | DMSO (pole 2) | 53 | one-way ANOVA with Dunnett's multiple comparisons test | F (4, 224)=11.47 | control | - |
|  | 5 nM (pole 2) | 44 |  |  | 0.1387 | NS |
|  | 10 nM (pole 2) | 44 |  |  | 0.7016 | NS |
|  | 15 nM (pole 2) | 44 |  |  | <0.0001 | **** |
|  | 20 nM (pole 2) | 44 |  |  | <0.0001 | **** |
| 4F | LoKI-off, pole 1 | 49 | unpaired two-tailed Student's t-test | t=4.770, df=96 | <0.0001 | **** |
|  | LoKI-off, pole 2 | 49 |  |  |  |  |
| 4F | LoKI-on, pole 1 | 49 | unpaired two-tailed Student's t-test | t=5.362, df=96 | <0.0001 | **** |
|  | LoKI-on, pole 2 | 49 |  |  |  |  |
| 4F | LoKI-off, pole 2 | 49 | unpaired two-tailed Student's t-test | t=3.382, df=96 | 0.0010 | ** |
|  | LoKI-on, pole 2 | 49 |  |  |  |  |
| 4H | DMSO | 76 | unpaired two-tailed Student's t-test | t=2.716, df=148 | 0.0074 | ** |
|  | 250 nM | 74 |  |  |  |  |
| S4B | DMSO, LoKI-off (pole 2) | 44 | unpaired two-tailed Student's t-test | t=1.163, df=102 | 0.2477 | NS |
|  | DMSO, LoKI-on (pole 2) | 60 |  |  |  |  |
| S4B | 100 nM, LoKI-off (pole 2) | 46 | unpaired two-tailed Student's t-test | t=0.09352, df=103 | 0.9257 | NS |
|  | 100 nM, LoKI-on (pole 2) | 59 |  |  |  |  |
| S4B | 500 nM, LoKI-off (pole 2) | 53 | unpaired two-tailed Student's t-test | t=2.612, df=111 | 0.0102 | * |
|  | 500 nM, LoKI-on (pole 2) | 60 |  |  |  |  |
| S4C | DMSO, LoKI-off (pole 2) | 52 | unpaired two-tailed Student's t-test | t=0.6178, df=109 | 0.5380 | NS |
|  | DMSO, LoKI-on (pole 2) | 59 |  |  |  |  |
| S4C | 100 nM, LoKI-off (pole 2) | 52 | unpaired two-tailed Student's t-test | t=0.1043, df=108 | 0.9172 | NS |
|  | 100 nM, LoKI-on (pole 2) | 58 |  |  |  |  |
| S4C | 250 nM, LoKI-off (pole 2) | 55 | unpaired two-tailed Student's t-test | t=0.7968, df=100 | 0.4275 | NS |
|  | 250 nM, LoKI-on (pole 2) | 47 |  |  |  |  |
| S4C | 500 nM, LoKI-off (pole 2) | 52 | unpaired two-tailed Student's t-test | t=0.2530, df=102 | 0.8007 | NS |
|  | 500 nM, LoKI-on (pole 2) | 52 |  |  |  |  |
| S4E | DMSO | 70 | unpaired two-tailed Student's t-test | t=2.106, df=135 | 0.0370 | * |
|  | 250 nM | 67 |  |  |  |  |
| 5C | Wildtype | 40 | unpaired two-tailed Student's t-test | t=2.140, df=72 | 0.0357 | * |
|  | Gravin -/- | 34 |  |  |  |  |
| 5E | Wildtype | 38 | unpaired two-tailed Student's t-test | t=2.538, df=78 | 0.0131 | * |
|  | Gravin -/- | 42 |  |  |  |  |

## Data availability

All raw image data can be provided upon request.

## Acknowledgments

We thank members of the Scott Lab for critical discussions and feedback on the manuscript, Patrina Pellett (GE Healthcare) for technical help with super-resolution imaging techniques, and Juan-Jesus Vicente (Wordeman Laboratory, UW) for help with experimental design, data analysis, and thoughtful discussion.

## Funding and additional information

This work was supported by NIH DK119186, DK119192 to J.D.S. Author contributions: P.J.B. and J.D.S. conceived of and designed the study. P.J.B. designed experiments. P.J.B., I.G. and R.M performed experiments. P.J.B. and I.G. analyzed the data. L.W. provided critical advice on experimental design, data analyses, and data interpretations. P.J.B. and I.G. designed and prepared figures. P.J.B. and J.D.S. wrote the manuscript.

## Conflict of interest

The authors declare no conflicts of interest in regards to this manuscript.

---

**Figure S1:**
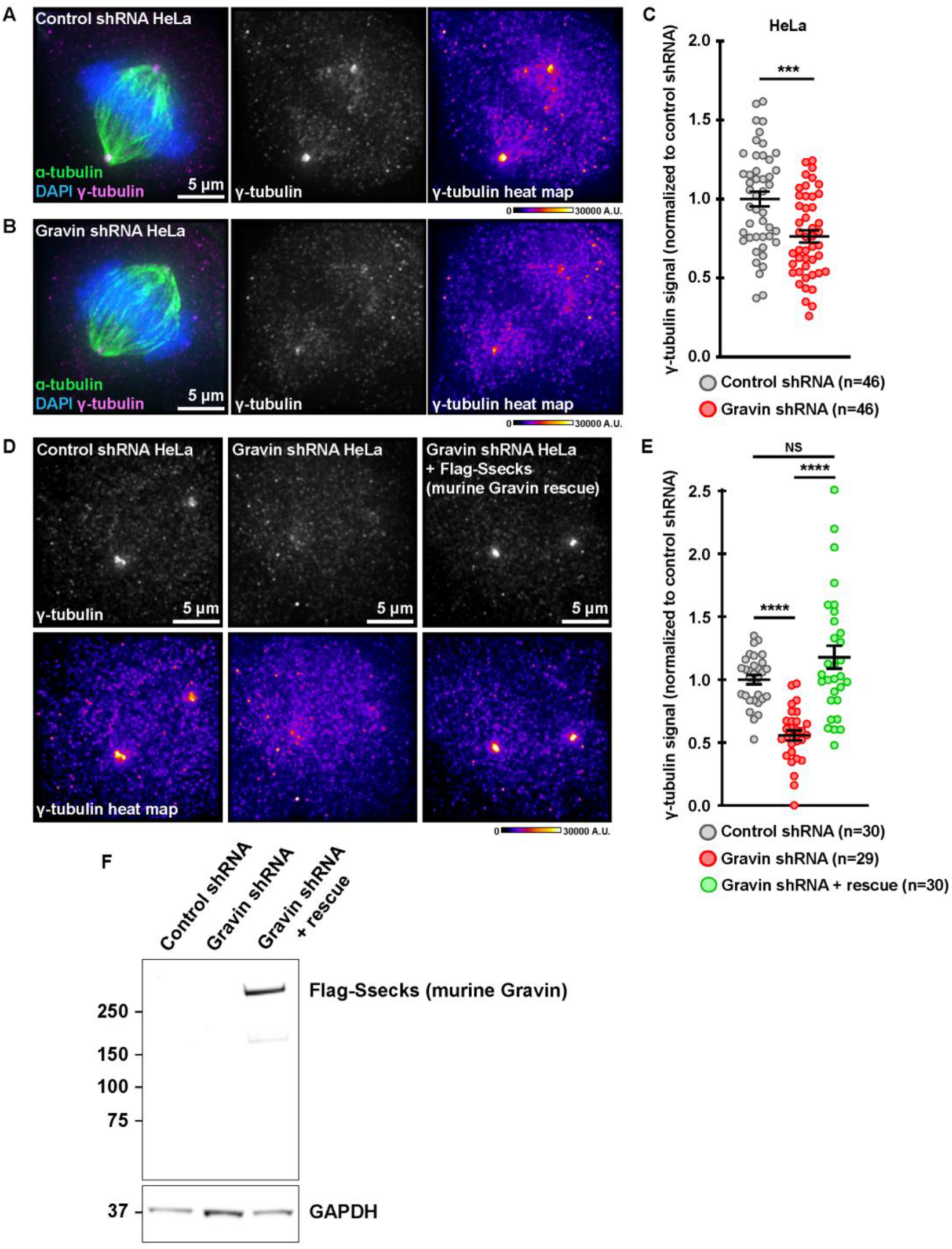
Loss of Gravin in HeLa cells perturbs accumulation of γ-tubulin at mitotic spindle poles. **A & B**. Immunofluorescence of representative control shRNA (**A**) and Gravin shRNA (**B**) mitotic HeLa cells. Composite images (left) show α-tubulin (green), DAPI (blue), and γ-tubulin (magenta). Distribution of γ-tubulin is represented in grayscale (middle) and with pseudo-color heat maps (right). Signal intensity scale (A.U.) is shown below. **C**. Quantification of amalgamated γ-tubulin immunofluorescence data at spindle poles in control shRNA (gray) and Gravin shRNA (red) HeLa cells; control shRNA, n=46, Gravin shRNA, n=46, ***p=0.0002. **D**. Immunofluorescence of representative mitotic HeLa cells stably expressing a control shRNA (left), Gravin shRNA (middle), or Gravin shRNA with Flag-Ssecks, a murine Gravin rescue construct (right). Distribution of γ-tubulin is represented in grayscale (top) and with pseudo-color heat maps (bottom). Signal intensity scale (A.U.) is shown below. **E**. Quantification of amalgamated γ-tubulin immunofluorescence data at spindle poles in control shRNA (gray), Gravin shRNA (red), and Gravin shRNA + rescue (green) HeLa cells. Prior to fixation, cells were treated for 24 hr with a CDK1 inhibitor, RO3306, followed by a 25 min washout to enrich for mitotic cells; control shRNA, n=30, Gravin shRNA, n=29, ****p<0.0001; Gravin shRNA, n=29, Gravin shRNA + rescue, n=30, ****p<0.0001; control shRNA, n=30, Gravin shRNA + rescue, n=30, p=0.0737. Points in **C** and **E** represent individual cells (n). Data in **C** and **E** are normalized to control shRNA. Cells were analyzed over three (**C**) or two (**E**) independent experiments (N=2-3). P values were calculated by unpaired two-tailed Student’s t-test. Data are mean ± s.e.m. NS, not significant. **F**. Immunoblot detection of Flag-Ssecks using an anti-Flag antibody (top) and GAPDH (bottom) in HeLa cells stably expressing a control shRNA (lane 1), Gravin shRNA (lane 2), or Gravin shRNA with the rescue construct (lane 3).

**Figure S2:**
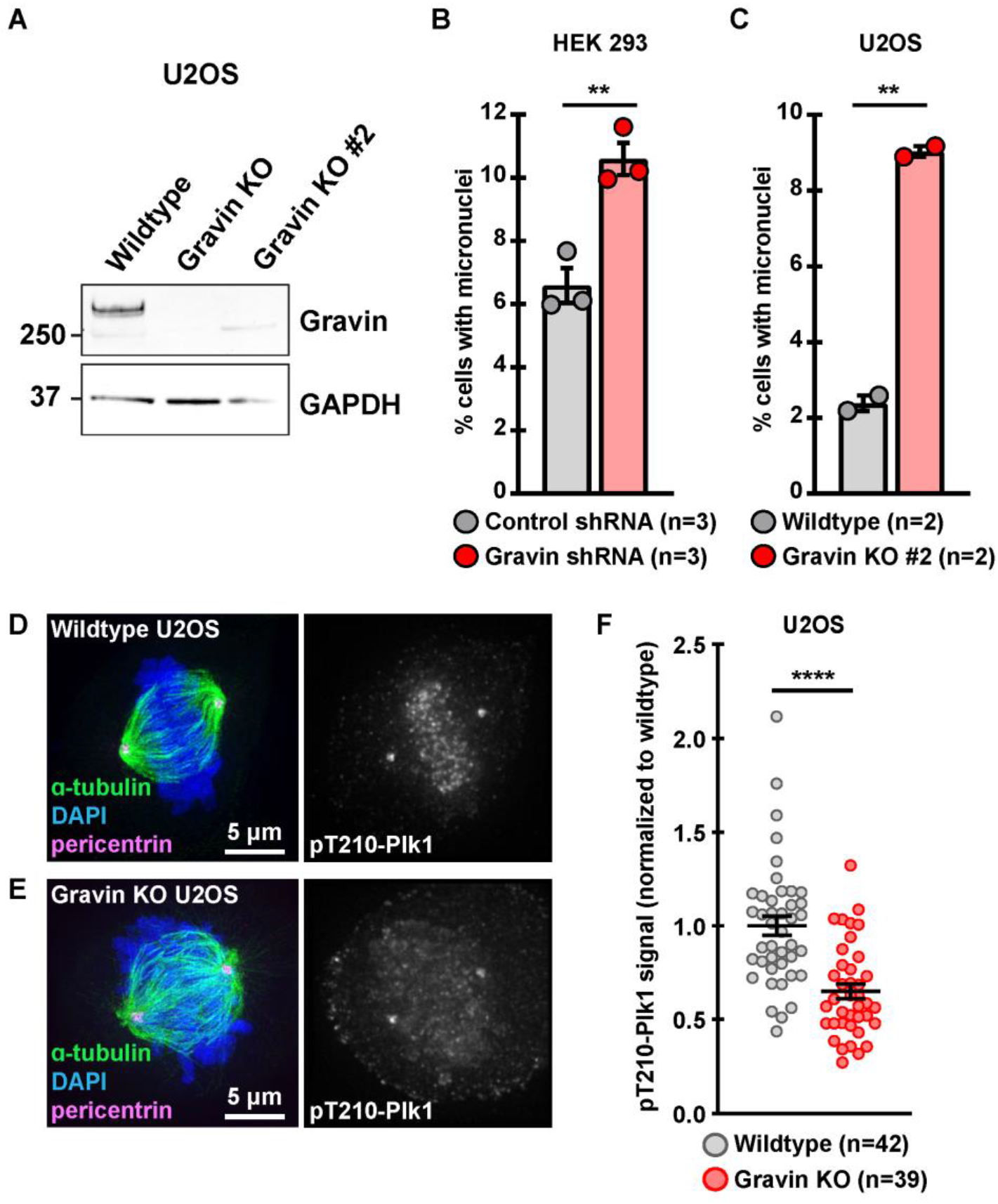
Further validation of Gravin knockout U2OS cells. **A**. Immunoblot detection of Gravin (top) using a rabbit antibody and GAPDH (bottom) in wildtype and Gravin KO clonal U2OS cells. **B & C**. Quantification representing the percent (%) of cells with micronuclei in HEK 293 (**B**) and wildtype and CRISPR/Cas9-edited U2OS clone 2 (**C**) cells. Points depict individual experiments (n); (**B**) control shRNA, n=3, Gravin shRNA, n=3, **p=0.0059; (**C**) wildtype, n=2, Gravin KO, n=2, **p=0.0014. A total of 1500 (**B**) or 1000 (**C**) cells were analyzed over three (**B**) or two (**C**) independent experiments. **D & E**. Immunofluorescence of representative wildtype (**D**) and Gravin KO (**E**) U2OS cells during mitosis. Composite SIM images (left) show α-tubulin (green), DAPI (blue), and pericentrin (magenta). Widefiled images show pT210-Plk1 signal in grayscale (right). **F**. Quantification of amalgamated pT210-Plk1 immunofluorescence data at spindle poles in wildtype (gray) and Gravin KO (red) U2OS cells. Points represent individual cells (n). Data are normalized to wildtype; wildtype, n=42, Gravin KO, n=39, ****p<0.0001; Immunofluorescence experiments were conducted three times (N=3). P values were calculated by unpaired two-tailed Student’s t-test. Data are mean ± s.e.m.

**Figure S3:**
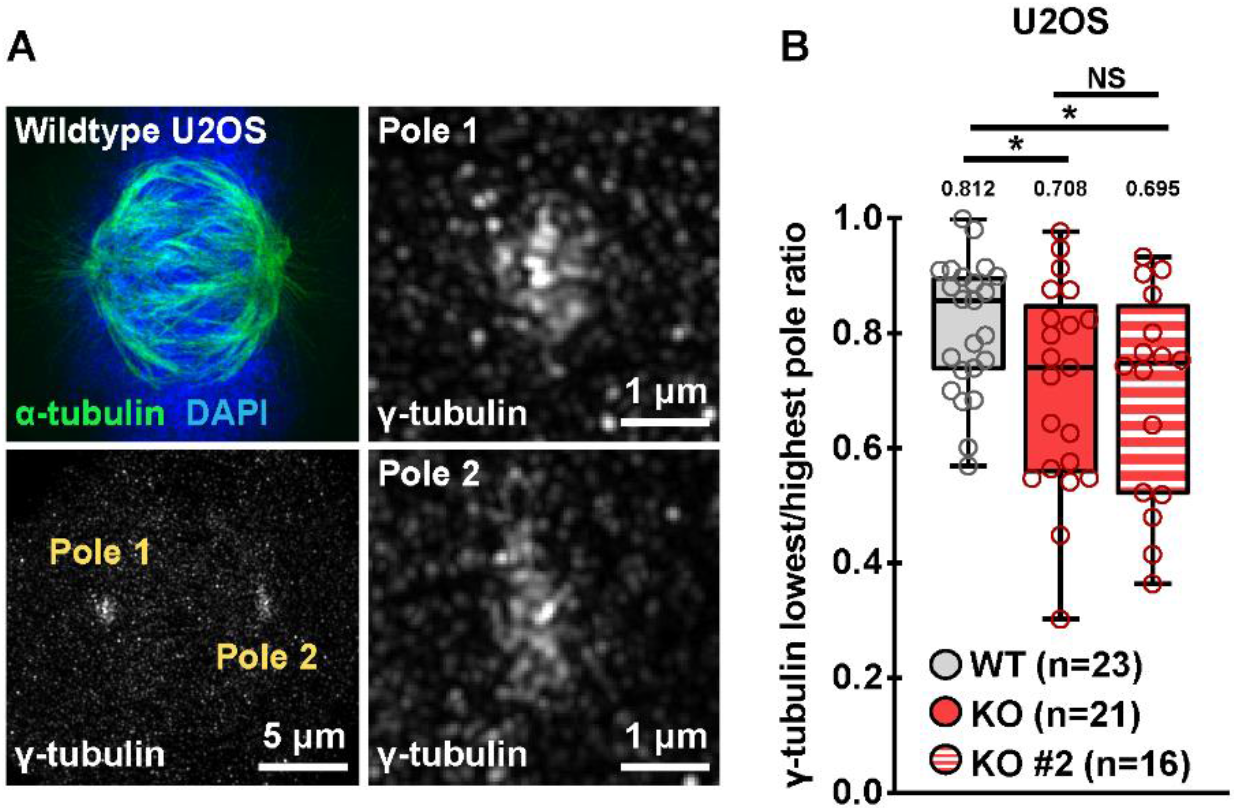
Pole-to-pole distribution of γ-tubulin in U2OS cells. **A**. SIM micrograph of a representative wildtype U2OS cell during mitosis. Composite image (top left) shows α-tubulin (green) and DAPI (blue). Grayscale image depicts γ-tubulin at both poles (bottom left). Magnified images reveal that pole 1 (top right) accumulates more γ-tubulin than pole 2 (bottom right). **B.** WT and Gravin KO conditions from Figure 4K with the addition of data for Gravin KO #2. Quantification of γ-tubulin immunofluorescence at each pole represented as box plots showing lowest/highest pole ratio in wildtype (gray), Gravin KO, (red) and Gravin KO #2 (red and white stripes) CRISPR/Cas9-edited U2OS mitotic cells. Mean values are indicated above each plot; WT, n=23, KO, n=21, *p=0.0263; WT, n=23, KO #2, n=16, *p=0.0190; KO, n=21, KO #2, n=16, p=0.8225. Points represent individual cells (n). Experiments were conducted three times (N=3) for WT and KO and two times (N=2) for KO #2. P values were calculated by unpaired two-tailed Student’s t-test. Data are mean ± s.e.m. NS, not significant.

**Figure S4:**
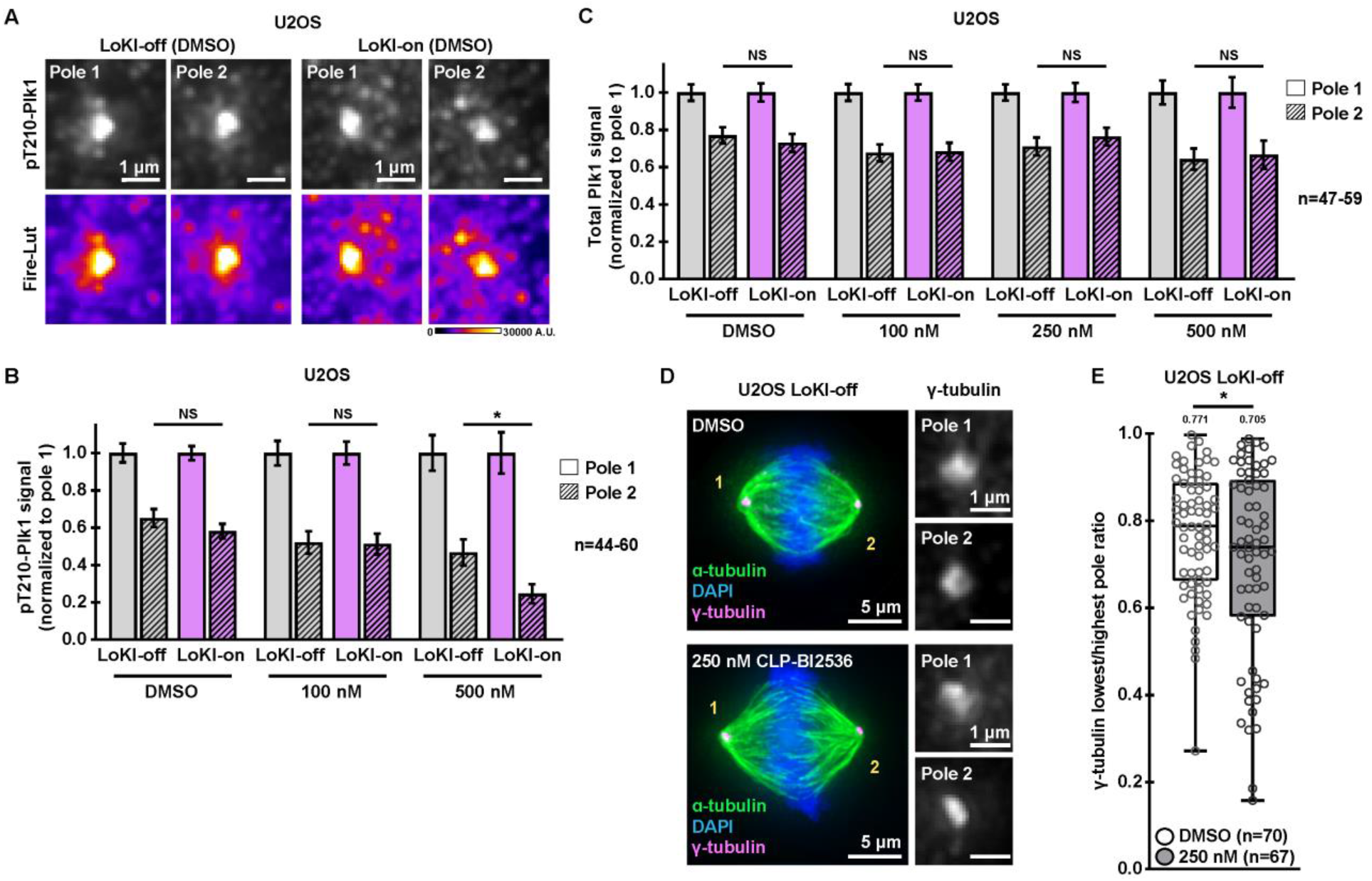
Plk1 inhibition promotes an asymmetric distribution of active kinase and γ-tubulin. **A**. Immunofluorescence of pT210-Plk1 at individual spindle poles in representative LoKI-off (left) and LoKI-on (right) U2OS cells treated with DMSO for 4 hr. Signal of active kinase is represented in grayscale (top) and with pseudo-color heat maps (bottom). Signal intensity scale (A.U.) is shown below. **B**. Quantification of pT210-Plk1 immunofluorescence at each pole after 4 hr treatment with DMSO or CLP-BI2536; pole 2: DMSO, LoKI-off, n=44, LoKI-on, n=60, p=0.2477; 100 nM, LoKI-off, n=46, LoKI-on, n=59, p=0.9257; 500 nM, LoKI-off, n=53, LoKI-on, n=60, *p=0.0102. **C**. Quantification of total Plk1 immunofluorescence at each pole after 4 hr treatment with DMSO or CLP-BI2536; pole 2: DMSO, LoKI-off, n=52, LoKI-on, n=59, p=0.5380; 100 nM, LoKI-off, n=52, LoKI-on, n=58, p=0.9172; 250 nM, LoKI-off, n=55, LoKI-on, n=47, p=0.4275; 500 nM, LoKI-off, n=52, LoKI-on, n=52, p=0.8007. **D**. Immunofluorescence of representative LoKI-off mitotic cells treated with DMSO (top) or 250 nM CLP-BI2536 (bottom) for 4 hr. Composite images (left) show α-tubulin (green), DAPI (blue), and γ-tubulin (magenta). Magnified grayscale images (right) show γ-tubulin signal at individual poles. **E**. Quantification of amalgamated γ-tubulin immunofluorescence data. Data represented as box plots showing lowest/highest pole ratio for LoKI-off cells after 4 hr treatment with DMSO (white) or 250 nM CLP-BI2536 (gray). Mean values are indicated above each plot; DMSO, n=70, 250 nM, n=67, *p=0.0370. Values depicted in **B**, **C** and **E** represent individual cells (n). Data in **B** and **C** are normalized to pole 1. Experiments were conducted at least three times (N=3). P values were calculated by unpaired two-tailed Student’s t-test. Data are mean ± s.e.m. NS, not significant.

## References

1. Scott, J. D., and Pawson, T. (2009) Cell signaling in space and time: where proteins come together and when they’re apart. Science 326, 1220–1224

2. Lemmon, M. A., Freed, D. M., Schlessinger, J., and Kiyatkin, A. (2016) The Dark Side of Cell Signaling: Positive Roles for Negative Regulators. Cell 164, 1172–1184

3. Langeberg, L. K., and Scott, J. D. (2015) Signalling scaffolds and local organization of cellular behaviour. Nat Rev Mol Cell Biol 16, 232–244

4. Gabrovsek, L., Bucko, P., Carnegie, G. K., and Scott, J. D. (2017) A-Kinase Anchoring Protein (AKAP). in Encyclopedia of Signaling Molecules (Choi, S. ed.), Springer New York, New York, NY. pp 1–6

5. Tasken, K., and Aandahl, E. M. (2004) Localized effects of cAMP mediated by distinct routes of protein kinase A. Physiol Rev 84, 137–167

6. Esseltine, J. L., and Scott, J. D. (2013) AKAP signaling complexes: pointing towards the next generation of therapeutic targets? Trends Pharmacol Sci 34, 648–655

7. Diviani, D., Langeberg, L. K., Doxsey, S. J., and Scott, J. D. (2000) Pericentrin anchors protein kinase A at the centrosome through a newly identified RII-binding domain. Curr Biol 10, 417–420

8. Chen, D., Purohit, A., Halilovic, E., Doxsey, S. J., and Newton, A. C. (2004) Centrosomal anchoring of protein kinase C betaII by pericentrin controls microtubule organization, spindle function, and cytokinesis. J Biol Chem 279, 4829–4839

9. Witczak, O., Skalhegg, B. S., Keryer, G., Bornens, M., Tasken, K., Jahnsen, T., and Orstavik, S. (1999) Cloning and characterization of a cDNA encoding an A-kinase anchoring protein located in the centrosome, AKAP450. EMBO J 18, 1858–1868

10. Canton, D. A., Keene, C. D., Swinney, K., Langeberg, L. K., Nguyen, V., Pelletier, L., Pawson, T., Wordeman, L., Stella, N., and Scott, J. D. (2012) Gravin is a transitory effector of polo-like kinase 1 during cell division. Mol Cell 48, 547–559

11. Hehnly, H., Canton, D., Bucko, P., Langeberg, L. K., Ogier, L., Gelman, I., Santana, L. F., Wordeman, L., and Scott, J. D. (2015) A mitotic kinase scaffold depleted in testicular seminomas impacts spindle orientation in germ line stem cells. Elife 4, e09384

12. Alvarado-Kristensson, M., Rodriguez, M. J., Silio, V., Valpuesta, J. M., and Carrera, A. C. (2009) SADB phosphorylation of gamma-tubulin regulates centrosome duplication. Nat Cell Biol 11, 1081–1092

13. Hendrickson, T. W., Yao, J., Bhadury, S., Corbett, A. H., and Joshi, H. C. (2001) Conditional mutations in gamma-tubulin reveal its involvement in chromosome segregation and cytokinesis. Mol Biol Cell 12, 2469–2481

14. Muller, H., Fogeron, M. L., Lehmann, V., Lehrach, H., and Lange, B. M. (2006) A centrosome-independent role for gamma-TuRC proteins in the spindle assembly checkpoint. Science 314, 654–657

15. Moritz, M., Braunfeld, M. B., Sedat, J. W., Alberts, B., and Agard, D. A. (1995) Microtubule nucleation by gamma-tubulin-containing rings in the centrosome. Nature 378, 638–640

16. Zheng, Y., Wong, M. L., Alberts, B., and Mitchison, T. (1995) Nucleation of microtubule assembly by a gamma-tubulin-containing ring complex. Nature 378, 578–583

17. Katsetos, C. D., Reddy, G., Draberova, E., Smejkalova, B., Del Valle, L., Ashraf, Q., Tadevosyan, A., Yelin, K., Maraziotis, T., Mishra, O. P., Mork, S., Legido, A., Nissanov, J., Baas, P. W., de Chadarevian, J. P., and Draber, P. (2006) Altered cellular distribution and subcellular sorting of gamma-tubulin in diffuse astrocytic gliomas and human glioblastoma cell lines. J Neuropathol Exp Neurol 65, 465–477

18. Caracciolo, V., D’Agostino, L., Draberova, E., Sladkova, V., Crozier-Fitzgerald, C., Agamanolis, D. P., de Chadarevian, J. P., Legido, A., Giordano, A., Draber, P., and Katsetos, C. D. (2010) Differential expression and cellular distribution of gamma-tubulin and betaIII-tubulin in medulloblastomas and human medulloblastoma cell lines. J Cell Physiol 223, 519–529

19. Cho, E. H., Whipple, R. A., Matrone, M. A., Balzer, E. M., and Martin, S. S. (2010) Delocalization of gamma-tubulin due to increased solubility in human breast cancer cell lines. Cancer Biol Ther 9, 66–76

20. Niu, Y., Liu, T., Tse, G. M., Sun, B., Niu, R., Li, H. M., Wang, H., Yang, Y., Ye, X., Wang, Y., Yu, Q., and Zhang, F. (2009) Increased expression of centrosomal alpha, gamma-tubulin in atypical ductal hyperplasia and carcinoma of the breast. Cancer Sci 100, 580–587

21. Maounis, N. F., Draberova, E., Mahera, E., Chorti, M., Caracciolo, V., Sulimenko, T., Riga, D., Trakas, N., Emmanouilidou, A., Giordano, A., Draber, P., and Katsetos, C. D. (2012) Overexpression of gamma-tubulin in non-small cell lung cancer. Histol Histopathol 27, 1183–1194

22. Friesen, D. E., Barakat, K. H., Semenchenko, V., Perez-Pineiro, R., Fenske, B. W., Mane, J., Wishart, D. S., and Tuszynski, J. A. (2012) Discovery of small molecule inhibitors that interact with gamma-tubulin. Chem Biol Drug Des 79, 639–652

23. Chinen, T., Liu, P., Shioda, S., Pagel, J., Cerikan, B., Lin, T. C., Gruss, O., Hayashi, Y., Takeno, H., Shima, T., Okada, Y., Hayakawa, I., Hayashi, Y., Kigoshi, H., Usui, T., and Schiebel, E. (2015) The gamma-tubulin-specific inhibitor gatastatin reveals temporal requirements of microtubule nucleation during the cell cycle. Nat Commun 6, 8722

24. Canton, D. A., and Scott, J. D. (2013) Anchoring proteins encounter mitotic kinases. Cell Cycle 12, 863–864

25. Colicino, E. G., Garrastegui, A. M., Freshour, J., Santra, P., Post, D. E., Kotula, L., and Hehnly, H. (2018) Gravin regulates centrosome function through PLK1. Mol Biol Cell 29, 532–541

26. Bucko, P. J., Lombard, C. K., Rathbun, L., Garcia, I., Bhat, A., Wordeman, L., Smith, F. D., Maly, D. J., Hehnly, H., and Scott, J. D. (2019) Subcellular drug targeting illuminates local kinase action. Elife 8

27. Zimmerman, W. C., Sillibourne, J., Rosa, J., and Doxsey, S. J. (2004) Mitosis-specific anchoring of gamma tubulin complexes by pericentrin controls spindle organization and mitotic entry. Mol Biol Cell 15, 3642–3657

28. Leibowitz, M. L., Zhang, C. Z., and Pellman, D. (2015) Chromothripsis: A New Mechanism for Rapid Karyotype Evolution. Annu Rev Genet 49, 183–211

29. Haren, L., Stearns, T., and Luders, J. (2009) Plk1-dependent recruitment of gamma-tubulin complexes to mitotic centrosomes involves multiple PCM components. PLoS One 4, e5976

30. Xu, D., and Dai, W. (2011) The function of mammalian Polo-like kinase 1 in microtubule nucleation. Proc Natl Acad Sci U S A 108, 11301–11302

31. Haren, L., Remy, M. H., Bazin, I., Callebaut, I., Wright, M., and Merdes, A. (2006) NEDD1-dependent recruitment of the gamma-tubulin ring complex to the centrosome is necessary for centriole duplication and spindle assembly. J Cell Biol 172, 505–515

32. Manning, J. A., Shalini, S., Risk, J. M., Day, C. L., and Kumar, S. (2010) A direct interaction with NEDD1 regulates gamma-tubulin recruitment to the centrosome. PLoS One 5, e9618

33. Zhang, X., Chen, Q., Feng, J., Hou, J., Yang, F., Liu, J., Jiang, Q., and Zhang, C. (2009) Sequential phosphorylation of Nedd1 by Cdk1 and Plk1 is required for targeting of the gammaTuRC to the centrosome. J Cell Sci 122, 2240–2251

34. Nauert, J. B., Klauck, T. M., Langeberg, L. K., and Scott, J. D. (1997) Gravin, an autoantigen recognized by serum from myasthenia gravis patients, is a kinase scaffold protein. Curr Biol 7, 52–62

35. Gelman, I. H. (2010) Emerging Roles for SSeCKS/Gravin/AKAP12 in the Control of Cell Proliferation, Cancer Malignancy, and Barriergenesis. Genes Cancer 1, 1147–1156

36. Parada, C. A., Osbun, J., Kaur, S., Yakkioui, Y., Shi, M., Pan, C., Busald, T., Karasozen, Y., Gonzalez-Cuyar, L. F., Rostomily, R., Zhang, J., and Ferreira, M., Jr. (2018) Kinome and phosphoproteome of high-grade meningiomas reveal AKAP12 as a central regulator of aggressiveness and its possible role in progression. Sci Rep 8, 2098

37. Steegmaier, M., Hoffmann, M., Baum, A., Lenart, P., Petronczki, M., Krssak, M., Gurtler, U., Garin-Chesa, P., Lieb, S., Quant, J., Grauert, M., Adolf, G. R., Kraut, N., Peters, J. M., and Rettig, W. J. (2007) BI 2536, a potent and selective inhibitor of polo-like kinase 1, inhibits tumor growth in vivo. Curr Biol 17, 316–322

38. Choi, M., Kim, W., Cheon, M. G., Lee, C. W., and Kim, J. E. (2015) Polo-like kinase 1 inhibitor BI2536 causes mitotic catastrophe following activation of the spindle assembly checkpoint in non-small cell lung cancer cells. Cancer Lett 357, 591–601

39. Lee, K. S., Burke, T. R., Jr., Park, J. E., Bang, J. K., and Lee, E. (2015) Recent Advances and New Strategies in Targeting Plk1 for Anticancer Therapy. Trends Pharmacol Sci 36, 858–877

40. Gutteridge, R. E., Ndiaye, M. A., Liu, X., and Ahmad, N. (2016) Plk1 Inhibitors in Cancer Therapy: From Laboratory to Clinics. Mol Cancer Ther 15, 1427–1435

41. Cheng, C. Y., Liu, C. J., Huang, Y. C., Wu, S. H., Fang, H. W., and Chen, Y. J. (2018) BI2536 induces mitotic catastrophe and radiosensitization in human oral cancer cells. Oncotarget 9, 21231–21243

42. Barr, F. A., Sillje, H. H., and Nigg, E. A. (2004) Polo-like kinases and the orchestration of cell division. Nat Rev Mol Cell Biol 5, 429–440

43. Lens, S. M., Voest, E. E., and Medema, R. H. (2010) Shared and separate functions of polo-like kinases and aurora kinases in cancer. Nat Rev Cancer 10, 825–841

44. Combes, G., Alharbi, I., Braga, L. G., and Elowe, S. (2017) Playing polo during mitosis: PLK1 takes the lead. Oncogene 36, 4819–4827

45. Lenart, P., Petronczki, M., Steegmaier, M., Di Fiore, B., Lipp, J. J., Hoffmann, M., Rettig, W. J., Kraut, N., and Peters, J. M. (2007) The small-molecule inhibitor BI 2536 reveals novel insights into mitotic roles of polo-like kinase 1. Curr Biol 17, 304–315

46. Klaeger, S., Heinzlmeir, S., Wilhelm, M., Polzer, H., Vick, B., Koenig, P. A., Reinecke, M., Ruprecht, B., Petzoldt, S., Meng, C., Zecha, J., Reiter, K., Qiao, H., Helm, D., Koch, H., Schoof, M., Canevari, G., Casale, E., Depaolini, S. R., Feuchtinger, A., Wu, Z., Schmidt, T., Rueckert, L., Becker, W., Huenges, J., Garz, A. K., Gohlke, B. O., Zolg, D. P., Kayser, G., Vooder, T., Preissner, R., Hahne, H., Tonisson, N., Kramer, K., Gotze, K., Bassermann, F., Schlegl, J., Ehrlich, H. C., Aiche, S., Walch, A., Greif, P. A., Schneider, S., Felder, E. R., Ruland, J., Medard, G., Jeremias, I., Spiekermann, K., and Kuster, B. (2017) The target landscape of clinical kinase drugs. Science 358

47. Kawashima, A. T., and Newton, A. C. (2020) Pharmacology on Target. Trends Pharmacol Sci 41, 227–230

48. Lane, H. A., and Nigg, E. A. (1996) Antibody microinjection reveals an essential role for human polo-like kinase 1 (Plk1) in the functional maturation of mitotic centrosomes. J Cell Biol 135, 1701–1713

49. Smith, F. D., Esseltine, J. L., Nygren, P. J., Veesler, D., Byrne, D. P., Vonderach, M., Strashnov, I., Eyers, C. E., Eyers, P. A., Langeberg, L. K., and Scott, J. D. (2017) Local protein kinase A action proceeds through intact holoenzymes. Science 356, 1288–1293

50. Smith, F. D., Reichow, S. L., Esseltine, J. L., Shi, D., Langeberg, L. K., Scott, J. D., and Gonen, T. (2013) Intrinsic disorder within an AKAP-protein kinase A complex guides local substrate phosphorylation. Elife 2, e01319

51. Nygren, P. J., Mehta, S., Schweppe, D. K., Langeberg, L. K., Whiting, J. L., Weisbrod, C. R., Bruce, J. E., Zhang, J., Veesler, D., and Scott, J. D. (2017) Intrinsic disorder within AKAP79 fine-tunes anchored phosphatase activity toward substrates and drug sensitivity. Elife 6

52. Smith, F. D., Omar, M. H., Nygren, P. J., Soughayer, J., Hoshi, N., Lau, H. T., Snyder, C. G., Branon, T. C., Ghosh, D., Langeberg, L. K., Ting, A. Y., Santana, L. F., Ong, S. E., Navedo, M. F., and Scott, J. D. (2018) Single nucleotide polymorphisms alter kinase anchoring and the subcellular targeting of A-kinase anchoring proteins. Proc Natl Acad Sci U S A 115, E11465–E11474

53. Fulcher, L. J., and Sapkota, G. P. (2020) Mitotic kinase anchoring proteins: the navigators of cell division. Cell Cycle 19, 505–524

54. Xia, W., Unger, P., Miller, L., Nelson, J., and Gelman, I. H. (2001) The Src-suppressed C kinase substrate, SSeCKS, is a potential metastasis inhibitor in prostate cancer. Cancer Res 61, 5644–5651

55. Bateman, N. W., Jaworski, E., Ao, W., Wang, G., Litzi, T., Dubil, E., Marcus, C., Conrads, K. A., Teng, P. N., Hood, B. L., Phippen, N. T., Vasicek, L. A., McGuire, W. P., Paz, K., Sidransky, D., Hamilton, C. A., Maxwell, G. L., Darcy, K. M., and Conrads, T. P. (2015) Elevated AKAP12 in paclitaxel-resistant serous ovarian cancer cells is prognostic and predictive of poor survival in patients. J Proteome Res 14, 1900–1910

